# Immunotherapy with immunocytokines and PD-1 blockade enhances the anticancer activity of small molecule-drug conjugates targeting carbonic anhydrase IX

**DOI:** 10.1101/2020.06.03.129049

**Authors:** Jacopo Millul, Christiane Krudewig, Aureliano Zana, Sheila Dakhel Plaza, Emanuele Puca, Alessandra Villa, Dario Neri, Samuele Cazzamalli

**Affiliations:** Philochem AG, Libernstrasse 3, CH-8112 Otelfingen (Switzerland); Laboratory for Animal Model Pathology, Universität Zürich, Winterthurerstrasse 268, CH-8057 Zurich (Switzerland); Department of Chemistry and Applied Biosciences, Swiss Federal Institute of Technology (ETH Zürich), Vladimir-Prelog-Weg 4, CH-8093 Zurich (Switzerland)

**Keywords:** Tumor Targeting, Small Molecule-Drug Conjugates, Immunotherapy, Carbonic Anhydrase IX, Therapy Studies

## Abstract

Small molecule-drug conjugates (SMDCs) represent an alternative to conventional antitumor chemotherapeutic agents, with the potential to improve the therapeutic window of cytotoxic payloads through active delivery at the site of the disease. In this article we describe novel combination therapies consisting of anti-Carbonic Anhydrase IX SMDCs combined with different immunomodulatory products. The therapeutic effect of the SMDCs was potentiated by combination with PD-1 blockade and with tumor-homing antibody-cytokine fusions in mouse models of renal cell carcinoma and colorectal cancer. The combination with L19-IL12, a fusion protein specific to the alternatively-spliced EDB domain of fibronectin containing the murine interleukin-12 moiety, was active also against large established tumors. Analysis of the microscopic structures of healthy organs performed three months after tumor eradication confirmed absence of pathological abnormalities in the healthy kidney, liver, lung, stomach and intestine. Our findings may be of clinical significance as they provide motivation for the development of combinations based on small molecule-drug conjugates and immunotherapy for the treatment of renal cell carcinoma and of hypoxic tumors.

## Introduction

The vast majority of cancer patients receive chemotherapy as part of combination treatments [1]. The clinical efficacy of conventional chemotherapeutic drugs is often limited by their unfavorable biodistribution profile [2]. The use of antibodies, specific to certain tumor-associated antigens and serving as drug delivery vehicles, may improve the therapeutic window of potent cytotoxic agents [3–5]. This approach has led to the marketing authorization of eight antibody-drug conjugates (ADCs; Mylotarg™, Kadcyla™, Adcetris™, Besponsa™, Polivy™, Padcev™, Enhertu™ and Trodelvy™) [4,6–9]. While the clinical benefit of ADCs has become evident, the therapeutic window of these biopharmaceuticals is narrower than what had initially been estimated on the basis of preclinical studies [10–12]. The toxicity profile of ADC products often involves organs which do not express the target antigen and may be substantial when approaching the maximal tolerated dose [13,14].

ADC products often extravasate slowly over the blood vessel walls and may exhibit a heterogeneous distribution within solid tumor masses [15,16]. Small molecule-based targeting agents may represent an alternative to antibodies. Excellent biodistribution profiles in mouse and man have been reported for small organic ligands specific to folate receptor [17], prostate-specific membrane antigen [18], fibroblast activation protein [19,20] and carbonic anhydrase IX [21]. The chemical conjugation of a small organic tumor targeting moiety to a cytotoxic drug leads to the generation of a novel class of therapeutic compounds named small molecule-drug conjugates (SMDCs) [22–24] which may represent an alternative to ADCs. Small ligands extravasate within seconds and may reach tumor cells far away from blood vessels [22]. In a recent direct comparison of an ADC and a SMDC directed against carbonic anhydrase IX, the small drug conjugate exhibited a better and more efficient tumor uptake, but both conjugates were efficacious in controlling tumor growth [25].

Carbonic anhydrase IX (CAIX) is a transmembrane homodimeric protein and a validated target for drug delivery applications [26]. CAIX is over-expressed in the majority of clear cell renal cell carcinomas (ccRCCs) and in hypoxic tumors, while its expression in normal tissues is mainly confined to certain structures of the gastrointestinal tract [27]. Both antibodies and small molecules have been successfully used to target CAIX-positive tumors *in vivo* [28–30]. Acetazolamide (AAZ), a small organic nanomolar binder of CAIX, represents a “portable” moiety for the active delivery of radionuclides, fluorophores and cytotoxic drugs [21,30–32].

We have recently described the generation of new anti-tumor SMDC products based on acetazolamide (AAZ) and on its affinity-matured derivative, called acetazolamide-plus (AAZ^+^) [33]. In this study, the two compounds were conjugated through the cleavable va-line-citrulline linker (Val-Cit) to the potent tubulin inhibitor monomethyl auristatin E (MMAE), also used in Adcetris® [34,35].

We observed a potentiation of anti-CAIX SMDC products using antibody-interleukin-2 and with antibody-interleukin-12 fusion proteins, that target the alternatively-spliced EDB domain of fibronectin [36]. Moreover, we studied the combination of SMDCs with a PD-1 immune checkpoint inhibitor [37]. All combinations were found to be safe over a prolonged period of time, as demonstrated by histopathological analysis performed on organs derived from cured animals. Our results provide a rationale for the clinical development of combination treatments based on the acetazolamide SMDC products, ideally in combination with tumor homing antibody-cytokine fusions or with anti-PD-1 antibodies.

## Materials and Methods

Detailed synthetic procedures and characterization of the presented compounds are described in the **Supplementary Information**.

### Preparation of SMDCs

AAZ-ValCit-MMAE was prepared following well-established synthetic procedures [31]. Briefly, commercially available MC-ValCit-PAB-MMAE (1 eq.) was reacted with compound **1** (4 eq. dissolved in 100 μl of DMF) in freshly degassed phosphate buffered saline (PBS; 50 mM phosphate, 100 mM NaCl, pH 7.4; 900 μl). The reaction was stirred at room temperature until completion (monitored by LC/MS). The crude was directly purified through RP-HPLC and solvents were removed by lyophilization. The identity and purity (98 %) of the final product was assessed by LC/MS.

AAZ^+^-ValCit-MMAE was prepared following well-established synthetic procedures [31]. Briefly, commercially available MC-ValCit-PAB-MMAE (1 eq.) was reacted with compound **2** (4 eq. dissolved in 100 μl of DMF) in freshly degassed phosphate buffered saline (PBS; 50 mM phosphate, 100 mM NaCl, pH 7.4; 900 μl). The reaction was stirred at room temperature until completion (monitored by LC/MS). The crude was directly purified through RP-HPLC and solvents were removed by lyophilization. The identity and purity (98 %) of the final product was assessed by LC/MS.

### Preparation of L19-IL12

The immunocytokine L19-IL12 was cloned as previously described [36]. Briefly, the fusion protein was produced in CHO-S mammalian cells and purified from the cell culture medium by affinity chromatography using a Protein A (Sino Biological) affinity column, as described previously [36]. After dialysis into PBS pH 7.4, the quality of the protein was assessed by size-exclusion chromatography and by SDS-PAGE [**Supplementary Figure 1**]. Binding to the cognate EDB antigen was assessed by Surface Plasmon Resonance [**Supplementary Figure 1**] following previously described procedures [36].

### Cell cultures

The human renal cell carcinoma cell line SKRC-52 was kindly provided by Professor E. Oosterwijk (Radbound University Nijmegen Medical Centre, Nijmegen, The Netherlands). SKRC-52 cells were cultured in RPMI medium (Invitrogen), supplemented with fetal calf serum (10%, FCS, Invitrogen) and Antibiotic-Antimycotic (1%, AA, Invitrogen) at 37°C, 5% CO_2_. Cells at 90-100% confluence were detached using Trypsin-EDTA 0.05% (Invitrogen) and re-seeded at a dilution of 1:6.

The murine colorectal carcinoma CT26-3E10 transfected cell [31] were cultured in RPMI medium (Invitrogen), supplemented with fetal calf serum (10%, FCS, Invitrogen) and Antibiotic-Antimycotic (1%, AA, Invitrogen) at 37°C, 5% CO_2_. Cells at 90-100% confluence were detached using Trypsin-EDTA 0.05% (Invitrogen) and re-seeded at a dilution of 1:6.

### Animal studies

All animal experiments were conducted in accordance with Swiss animal welfare laws and regulations under the license number ZH004/18 granted by the Veterinäramt des Kantons Zürich.

### Implantation of subcutaneous tumors

SKRC-52 cells were grown to 80-100% confluence and detached with Trypsin-EDTA 0.05% (Invitrogen). Cells were washed once with Hank’s Balanced Salt Solution (HBSS, Thermo Fisher Scientific, pH 7.4), counted and re-suspended in HBSS. Aliquots of 5-10 × 10^6^ cells were resuspended in 150 μl of HBSS and injected subcutaneously in the right flank of female athymic BALB/c *nu/nu* mice (8-10 weeks of age, Janvier).

CT26-3E10 cells were grown to 80-100% confluence and detached with Trypsin-EDTA 0.05% (Invitrogen). Cells were washed once with Hank’s Balanced Salt Solution (HBSS, Thermo Fisher Scientific, pH 7.4), counted and re-suspended in HBSS. Aliquots of 6 × 10^6^ cells were resuspended in 150 μl of HBSS and injected subcutaneously in the right flank of female athymic BALB/c *nu/nu* mice (8-10 weeks of age, Janvier).

### Quantification of MMAE in SKRC-52 tumors

SKRC-52 tumor cells were implanted into female BALB/c athymic *nu/nu* mice and allowed to grow to an average volume of 100 mm^3^. AAZ-ValCit-MMAE was injected through tail vein injection (250 nmol/Kg) and animals were euthanized at different time points (i.e., 10 min, 1h, 3h, 6h and 24h after the administration). The tumors were collected, frozen and stored at −80°C prior to sample preparation and LC/MS quantification of MMAE. Blood was collected and incubated for 20 min in Microtainer tubes containing lithium heparin (BD Microtainer Tube). Plasma was obtained by centrifugation for 15 min at 3’000 rpm using a refrigerated centrifuge and stored at −80°C prior to sample preparation and LC/MS quantification of MMAE. The detailed protocol for processing the samples is described in the **Supplementary information.**

### Dose escalation study with L19-IL12

A dose escalation study with L19-IL12 was performed in athymic BALB/c *nu/nu* mice (8-10 weeks of age, Janvier). Four different doses of the immunocytokine were tested following the schedule reported in the **Supplementary Information** (three intravenous injections per animal at 0.6, 0.8, 1 or 1.2 mg/kg). Animals were weighed daily to assess acute toxicity of the immunocytokine at the different doses.

### Therapy experiments

SKRC-52 or CT26-3E10 tumor cells were implanted into female BALB/c *nu/nu* mice and allowed to grow to an average volume of 100 mm^3^ or 200 mm^3^. Mice were randomly assigned into therapy groups of 3, 4 or 5 animals.

Intravenous injections of AAZ-ValCit-MMAE (250 nmol/kg), AAZ^+^-ValCit-MMAE (250 nmol/kg), L19-IL2 (2.5 mg/kg), anti-PD-1 (10 mg/kg), L19-IL12 (1.2 mg/kg) or vehicle were performed with the schedules indicated in the text and in Fig. 2, 3 and 4. Compounds **1** and **2** were injected as sterile PBS solution with 1% of DMSO. L19-IL12 and anti-PD-1 were injected as sterile PBS solution. L19-IL2 was injected in sterile formulation buffer (Philogen).

Tumors were measured with an electronic caliper and animals were weighted daily. Tumor volume (mm^3^) was calculated with the formula (long side, mm) x (short side, mm) x (short side, mm) x 0.5. Animals were sacrificed when one or more termination criteria indicated by the experimental license were reached (e.g. weight loss > 15%).

Prism 6 software (GraphPad Software) was used for data analysis (regular two-way ANOVA followed by Bonferroni test).

### Evaluation of chronic toxicity by plasma analysis and *ex vivo* histology

SKRC-52 bearing mice treated with vehicle, SMDC products and immunocytokines as single agents or their combinations (n = 3 per group) were sacrificed 90 days after treatment. The plasma was collected for each mouse and analyzed to detect the level of creatinine and urea. A complete gross post mortem examination had been performed on each mouse, gross abnormalities recorded and all tissues collected in 10% buffered formalin for histological analysis. The tissue was trimmed, dehydrated, and embedded in paraffin wax. Sections of 3-4 μm thickness were prepared, mounted on glass slides, deparaffinized in xylene, and rehydrated through graded alcohols, before staining with hematoxylin and eosin (HE) for the histological examination. Samples containing bone (bone marrow) were demineralized after fixation with EDTA for 48 hours prior to further processing as described above.

The microscopic morphology of different organs (kidney, liver, lung, stomach, bone marrow and spleen) was blindly analyzed by a board-certified pathologist. Separately and afterwards they were compared to corresponding control samples collected from a nontreated animal to highlight structural changes deriving from the different treatments.

## Results

### Production and characterization of SMDCs and immunocytokines

The structure of AAZ-ValCit-MMAE and AAZ^+^-ValCit-MMAE are shown in **Figure 1A**. The peptide precursor compounds **1** and **2**, featuring respectively AAZ or AAZ^+^ as CAIX targeting moiety [31], were prepared by solid phase peptide synthesis following a conventional Fmoc-strategy and purified by reverse phase HPLC. Compounds **1** and **2** were coupled in solution to the linker-payload module used for Adcetris™, composed by the cleavable linker Valine-Citrulline, a self-immolative para-aminobenzyl carbamate moiety and the potent anti-tubulin poison monomethyl auristatin E (MMAE). The resulting AAZ-ValCit-MMAE and AAZ^+^-ValCit-MMAE products were used for therapy studies in tumorbearing mice.

**Figure 1:**
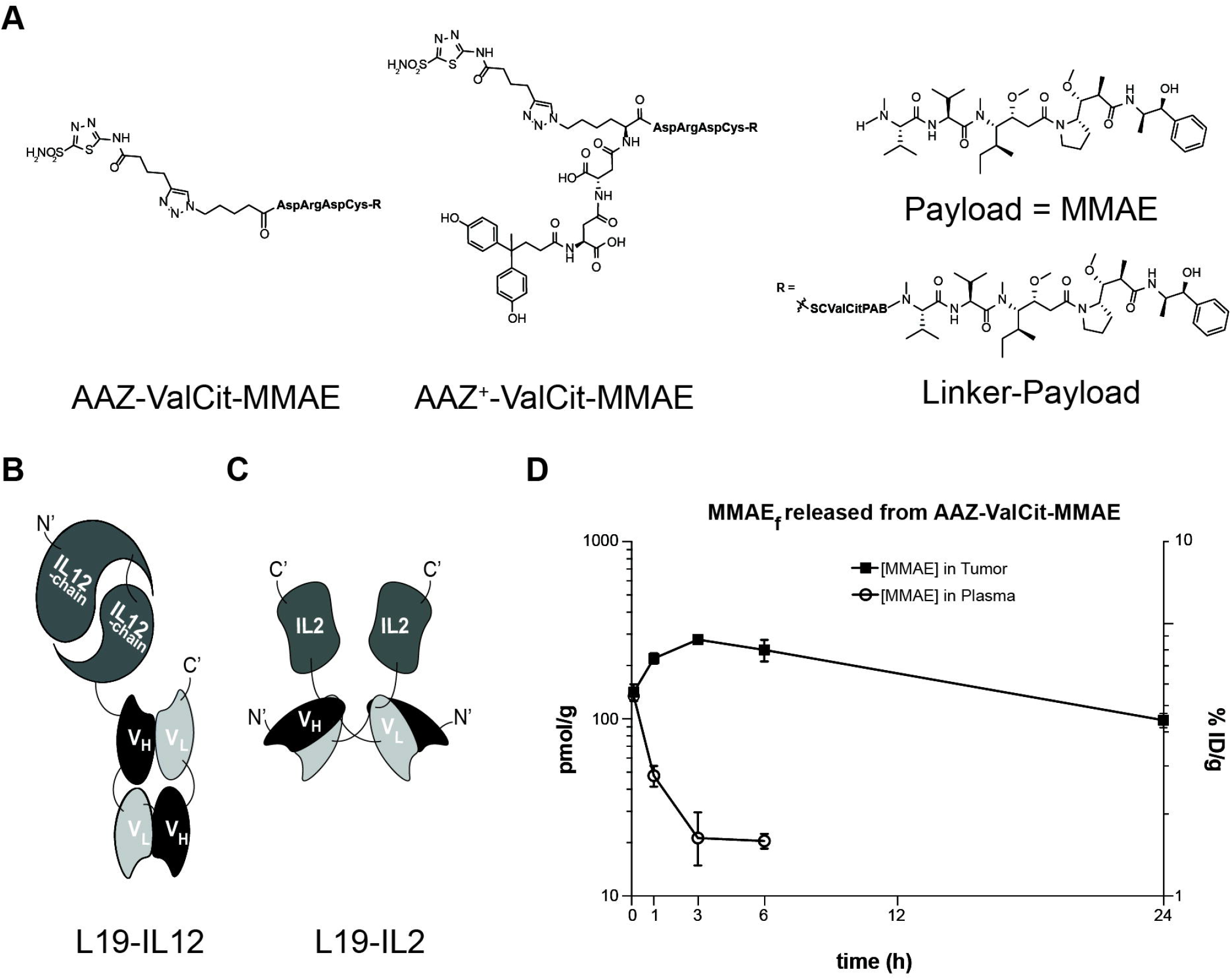
Small molecule-drug conjugates targeting CAIX, antibody-fusion proteins targeting EDB domain of Fibronectin and bioanalysis of MMAE released *in vivo* by AAZ-ValCit-MMAE. (**A**) Chemical structures of AAZ-ValCit-MMAE, AAZ^+^-ValCit-MMAE and of corresponding payload and linker-payload modules. (**B**) Schematic structure of L19-IL12 (**C**) and of L19-IL2. (**D**) Quantification of free MMAE in SKRC-52 tumors and plasma at different time points after intravenous administration of AAZ-ValCit-MMAE in tumor bearing mice at a dose of 250 nmol/Kg. SC = succinimido-caproyl; PAB = para-amino-benzyl.

The immunocytokine L19-IL12, recently described by our group, is a fusion protein consisting of the L19 antibody in tandem diabody format fused to a single-chain of the murine IL-12 by the (GGGGS)3 15-amino-acid linker [36] [**Figure 1B**]. We have also previously reported the fusion of the L19 antibody with interleukin-2 [31] [**Figure 1C**]. Structural and analytical information for all products is reported in the **Supplementary Information and Supplementary Figure 1.**

### Quantification study of MMAE in SKRC-52 tumor

We quantified the amount of the MMAE cytotoxic payload released in tumors and in plasma after injection of a single dose of AAZ-ValCit-MMAE pro-drug (250 nmol/Kg) in SKRC-52 tumor bearing mice [**Figure 1D**]. A high and stable accumulation of free MMAE was observed in tumor samples. The highest concentration was obtained in tumor samples collected 3 hours after the intravenous injection, with values corresponding to 270 pmol/gram of tissue (i.e., to 5.4% of the injected dose per gram). Free MMAE levels in circulation quickly went down to values lower that 1% of the injected dose per mL of plasma already 1 hour after injection of the AAZ-ValCit-MMAE prodrug.

### Therapy experiment of combination of SMDCs and L19-IL2 in SKRC-52 renal cell carcinoma model

We had previously reported that L19-IL2 potentiates the anti-cancer activity of AAZ^+^-ValCit-MMAE in two mouse models of cancer [31] and that the AAZ^+^ moiety targets tumors more efficiently than AAZ [38]. Here, we performed a comparison of AAZ-ValCit-MMAE and AAZ^+^-ValCit-MMAE in combination with L19-IL2, in order to study the activity and tolerability of the two drugs, AAZ-ValCit-MMAE and AAZ^+^-ValCit-MMAE were administered at 250 nmol/Kg, a dose that was previously identified as safe and effective in the SKRC-52 renal cell carcinoma model, [21,25,31]. The antibody-fusion protein L19-IL2 was administered at the dose of 2.5 mg/Kg, as previously described [31]. Therapy started when the average volume of established tumors reached 100 mm^3^. We compared the simultaneous administration of SMDC and L19-IL2 with a modified schedule, featuring a 24-hour delay between the two injections [**Figure 2A and 2B**], as it has previously been suggested that immunotherapy may benefit from the damage caused to the tumor by cytotoxic agents [39]. A potent *in vivo* anti-cancer activity of both SMDCs in combination with L19-IL2 was observed in both schedules [**Figure 2A and 2B**]. AAZ^+^-ValCit-MMAE was more active in this study, leading to 4/4 cures in combination with L19-IL2 in the sequential administration schedule and 2/4 cures upon simultaneous administration [**Figure 2B**]. All treatments were well tolerated at the administered doses [**Figure 2C and 2D**].

**Figure 2:**
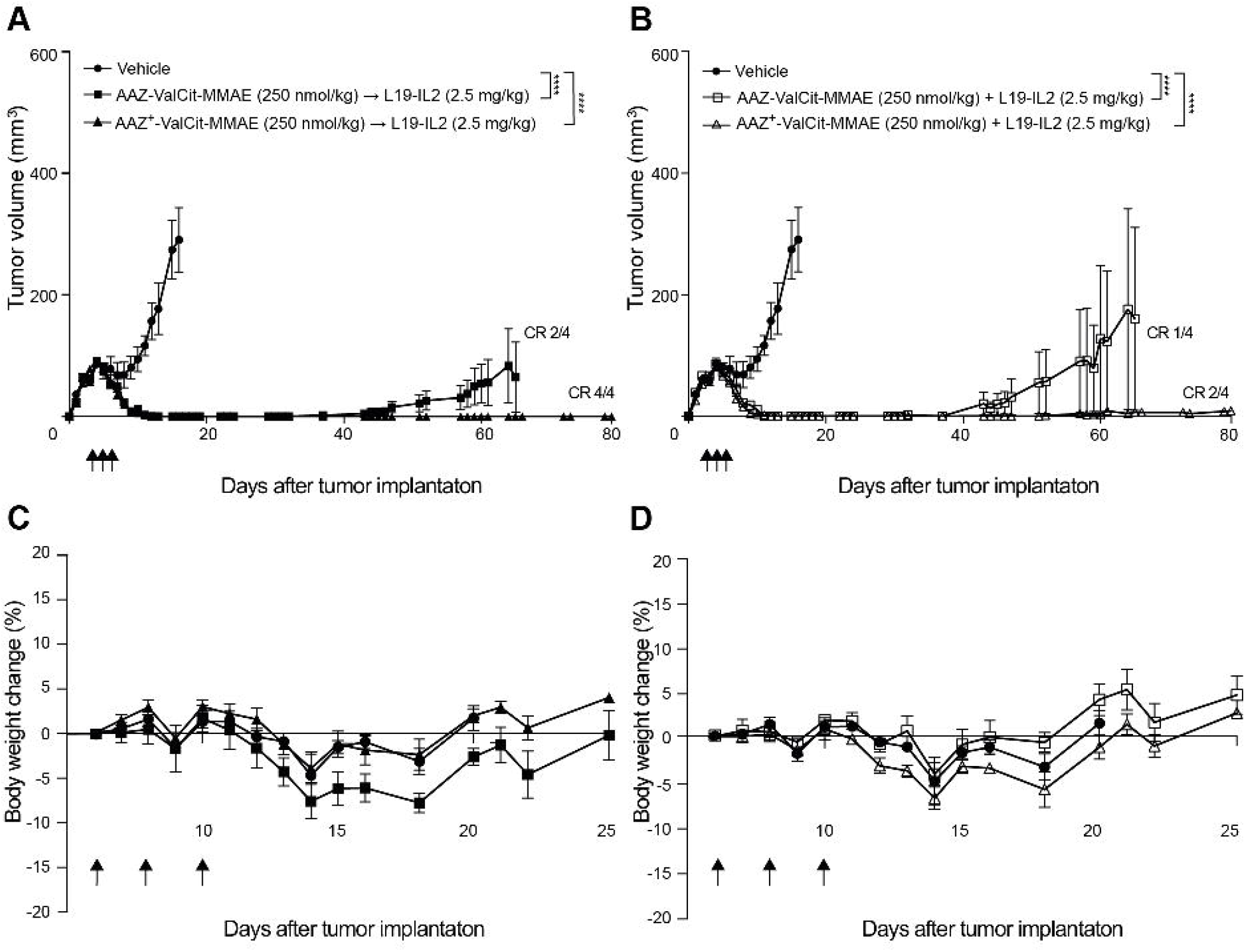
*In vivo* efficacy of AAZ-ValCit-MMAE and AAZ^+^-ValCit-MMAE in combination with L19-IL2 in BALB/c *nu/nu* mice bearing established subcutaneous SKRC-52 renal cell carcinoma. (**A**) Therapeutic activity of the different treatments as assessed by measurement of tumor volume (mm^3^) during therapy experiment. Data points represent mean tumor volume ± SEM (n = 4 per group). The two treatments were administered intravenously as bolus injections with a 24-hour gap, starting from the SMDCs. (**B**) Therapeutic activity of the different treatments given simultaneously, without time gap between the two injections. In both cases, the combination of the AAZ^+^-ValCit-MMAE with L19-IL2 exhibited a superior anti-tumor activity when compared with vehicle and with the combination of AAZ-ValCit-MMAE with L19-IL2. (**C**) Tolerability of the different treatments as assessed by the evaluation of changes (%) in body weight during the experiment when the two treatments were given with a 24 hours gap, starting from the SMDCs. (**D**) Tolerability of the different treatments as assessed by the evaluation of changes (%) in body weight during the experiment when the two treatments were given simultaneously. All treatments were well tolerated and did not have an impact on the body weight of the animals. **** indicates p<0.0001; ** indicates p<0.01; * indicates p<0.05; ns indicates p>0.05 (2-way ANOVA test, followed by Bonferroni post-test).

### Therapy experiment of combination of SMDCs and anti-PD-1 in CT26-3E10 colorectal cell carcinoma model

The anti-cancer activity of AAZ-ValCit-MMAE, of AAZ^+^-ValCit-MMAE, of anti-PD-1 and of their combination was assessed in BALB/c mice bearing subcutaneously CT26-3E10 colorectal cell carcinoma. AAZ-ValCit-MMAE and AAZ^+^-ValCit-MMAE were administered at 250 nmol/Kg, as previously mentioned. The anti-PD-1 antibody was administered at the dose of 10 mg/Kg, as previously described [37]. Therapy started when the average volume of established tumors had reached 100 mm^3^. The compounds were administered the same day, with a 6 hours interval between the two injections, starting with the SMDC [**Figure 3A**]. A potent anti-cancer activity of AAZ-ValCit-MMAE, AAZ^+^-ValCit-MMAE and anti-PD-1 were observed *in vivo* when the products were given as combination, while the single treatment with anti-PD-1 was ineffective [**Figure 3A**]. All treatments were well tolerated at the doses administered [**Figure 3B**].

**Figure 3:**
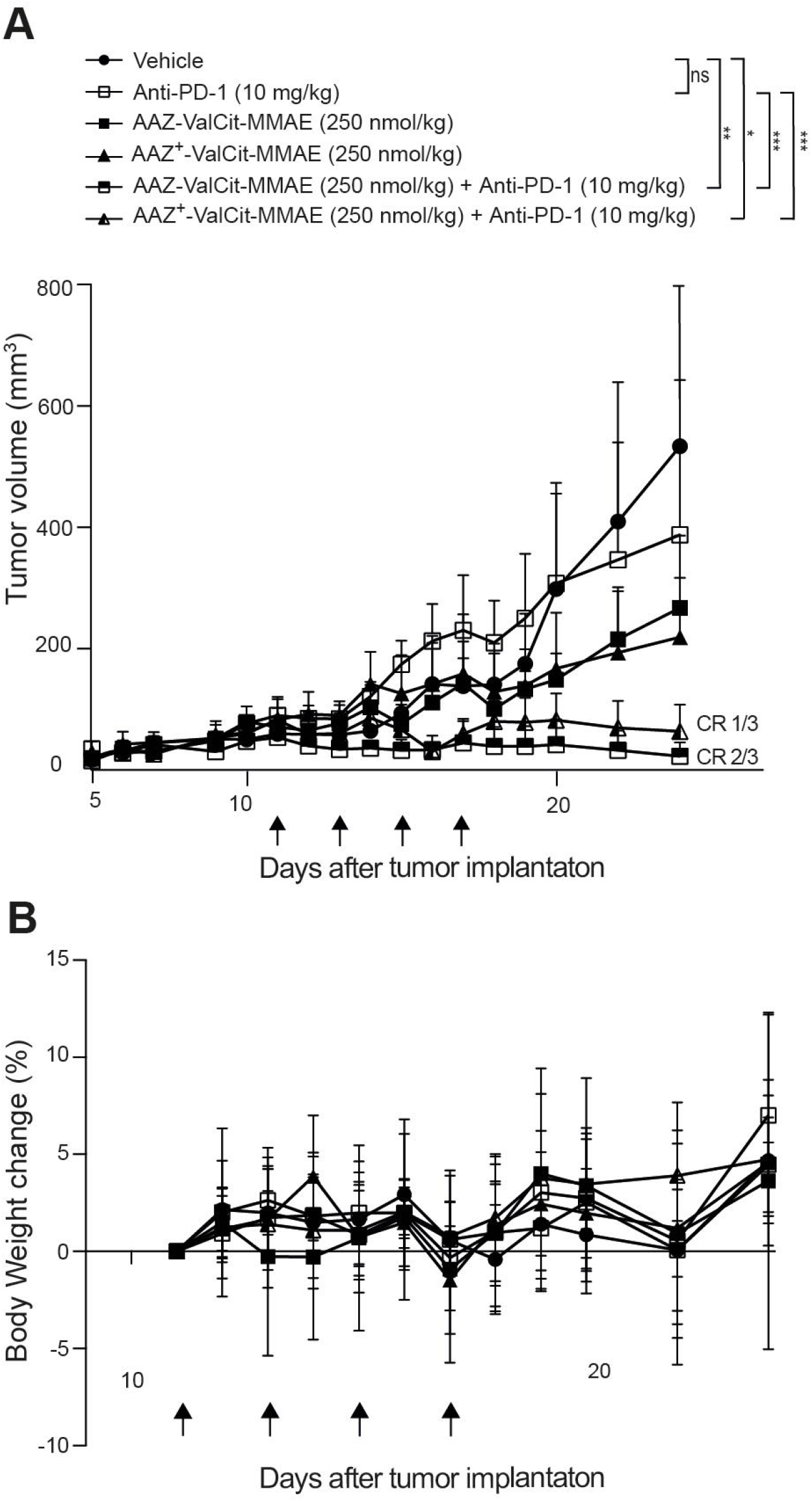
*In vivo* efficacy of AAZ-ValCit-MMAE and AAZ^+^-ValCit-MMAE in combination with Anti-PD-1 in BALB/c mice bearing established subcutaneous CT26.3E10 colorectal cell carcinoma. (**A**) Therapeutic activity of the different treatments as assessed by measurement of tumor volume (mm^3^) during therapy experiment. Data points represent mean tumor volume ± SEM (n = 3 per group). The two treatments were administered intravenously as bolus injections with a 6 hours window, starting from the SMDCs. The combination of the SMDCs with anti-PD-1 exhibited a superior anti-tumor activity when compared with vehicle, AAZ-ValCit-MMAE, AAZ^+^-ValCit-MMAE or anti-PD-1 only. (**B**) Tolerability of the different treatments as assessed by the evaluation of changes (%) in body weight during the experiment. All treatments were well tolerated and did not have an impact on the body weight of the animals. **** indicates p<0.0001; ** indicates p<0.01; * indicates p<0.05; ns indicates p>0.05 (2-way ANOVA test, followed by Bonferroni post-test).

### Therapy experiments of combination of SMDC and L19-IL12 in SKRC-52 renal cell carcinoma model

The anti-cancer activity of AAZ^+^-ValCit-MMAE, of L19-IL12 and of their combination was assessed in BALB/c nude mice bearing subcutaneously-grafted SKRC-52 renal cell carcinoma. AAZ^+^-ValCit-MMAE was administered at 250 nmol/Kg. The antibody-fusion protein L19-IL12 was administered at the dose of 1.2 mg/Kg. This dose was considered to be safe based on a preliminary dose escalation experiment in nude mice [**Supplementary Information**]. Therapy started when the average volume of established tumors reached 100 mm^3^. The two compounds were administered according to the schedule described in **Figure 4A**. A potent anti-cancer activity for AAZ^+^-ValCit-MMAE and for L19-IL12 was observed *in vivo* when products were given as monotherapy. Moreover, all animals treated with the combination of the two products experienced a complete and durable tumor eradication [**Figure 4A**]. AAZ^+^-ValCit-MMAE, L19-IL12 and their combination treatment were well tolerated at the administered doses [**Figure 4B**]. Cured animals were kept alive and tumor-free for 90 days after the administration of the first dose of SMDC [**Figure 4A**].

**Figure 4:**
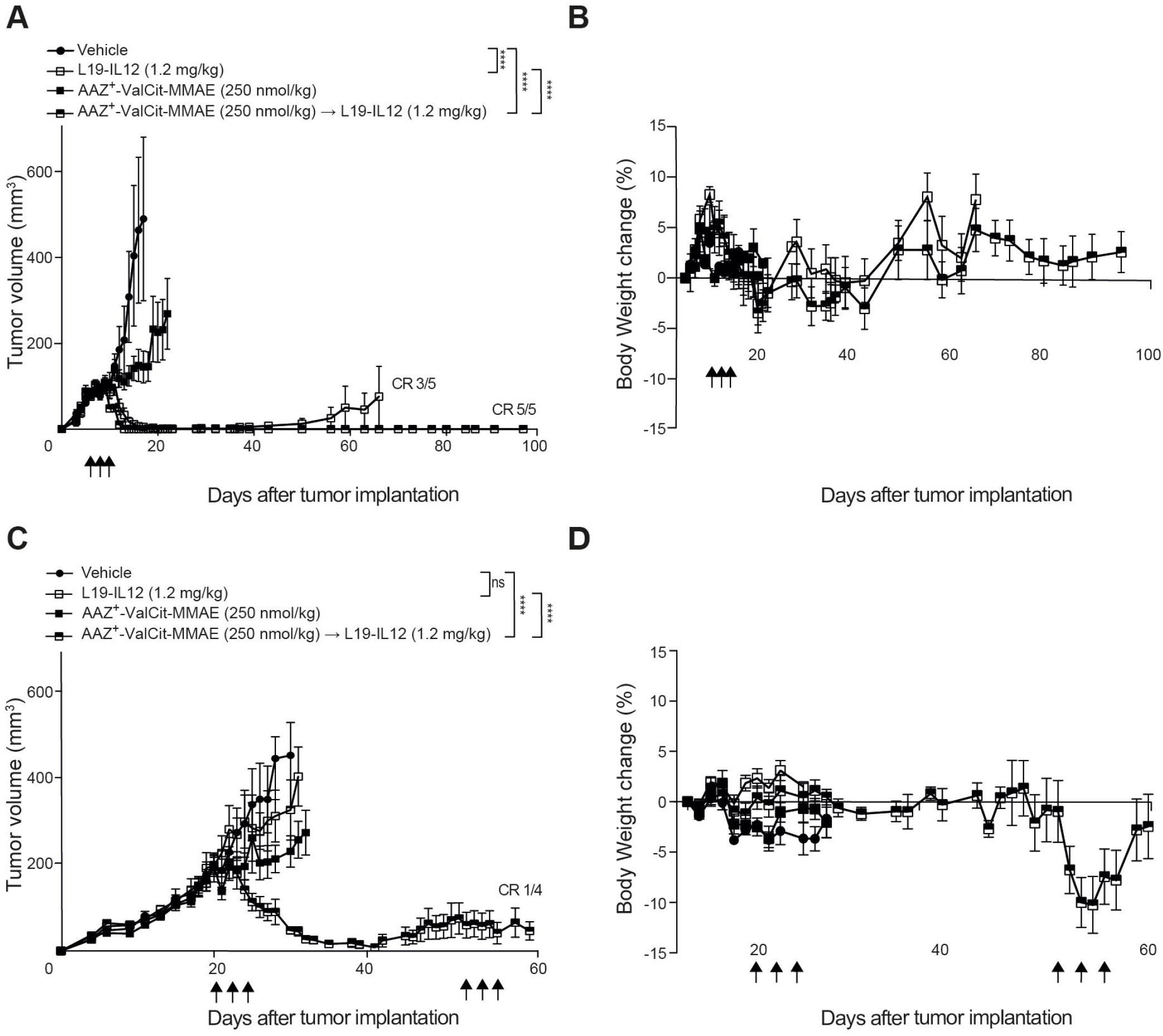
*In vivo* efficacy of AAZ^+^-ValCit-MMAE in combination with L19-IL12 in BALB/c *nu/nu* mice bearing established subcutaneous SKRC-52 renal cell carcinoma with different initial tumor burden. (**A**) Therapeutic activity of the different treatments as assessed by measurement of tumor volume (mm^3^) during therapy experiment. Therapy experiment started when the average tumor volume was about 100 mm^3^. Data points represent mean tumor volume ± SEM (n = 5 per group). The two treatments were administered intravenously as bolus injections with a 24-hour gap, starting from the SMDCs. The combination of AAZ^+^-ValCit-MMAE with L19-IL12 exhibited a superior anti-tumor activity when compared with vehicle, L19-IL12 or AAZ^+^-ValCit-MMAE only. (**B**) Tolerability of the different treatments as assessed by the evaluation of changes (%) in body weight during the experiment. All treatments were well tolerated and did not have an impact on the body weight of the animals. **** indicates p<0.0001; ** indicates p<0.01; * indicates p<0.05; ns indicates p>0.05 (2-way ANOVA test, followed by Bonferroni post-test). (**C**) Therapeutic activity of AAZ^+^-ValCit-MMAE, L19-IL12 as single agents or in combination as assessed by measurement of tumor volume (mm^3^) during therapy experiment. Therapy experiment started when the average tumor volume was about 200 mm^3^. Data points represent mean tumor volume ± SEM (n = 4 per group). The two treatments were administered intravenously as bolus injections with a 24-hour gap, starting from the SMDCs. (**D**) Tolerability of the different treatments as assessed by the evaluation of changes (%) in body weight during the experiment. **** indicates p<0.0001; ** indicates p<0.01; * indicates p<0.05; ns indicates p>0.05 (2-way ANOVA test, followed by Bonferroni post-test).

In a second therapy experiment, we tested the anti-cancer activity of the combination AAZ^+^-ValCit-MMAE and L19-IL12 on larger established SKRC-52 tumors (average volume of 200 mm^3^) implanted in BALB/c nude mice. The administration of AAZ^+^-ValCit-MMAE and L19-IL12 as single agents resulted in a partial tumor growth retardation, compared to saline-treated animals used as controls [**Figure 4C**]. Interestingly, only the combination of AAZ^+^-ValCit-MMAE and L19-IL12 induced tumor regression in this setting. All treatments were well tolerated and no sign of acute toxicity was observed [**Figure 4D**]. Mice in the combination group received a second cycle of injections (i.e. AAZ^+^-ValCit-MMAE and L19-IL12) on therapy day 52. During this second round of treatments, an acute and reversible body weight loss was observed as sign of toxicity [**Figure 4D**]. Mice treated with the combination were kept alive for more than 60 days [**Figure 4C**].

### Analysis of chronic toxicity of combination treatment of SMDCs with L19-IL2

Gross examination of mice revealed a slightly to moderately enlarged spleen of all animals treated with L19-IL2 alone or in combination with SMDCs compared to the control group. Histologically, none of the treatments caused cellular infiltration and/or connective tissue (fibrosis) in liver and kidney of the treated animal in comparison with specimens from control animals. No signs of toxicity were found in other healthy organs, including lung and stomach that are known to be exposed to higher concentration of AAZ-based derivatives [**Figure 5A**] [30]. Urea and creatinine plasma levels measured three months after treatment with SMDCs, L19-IL2 and combinations (blood chemistry for kidney chronic toxicity) revealed no significant differences between treatment groups and compared to the control group [**Figure 5B and 5C**].

**Figure 5:**
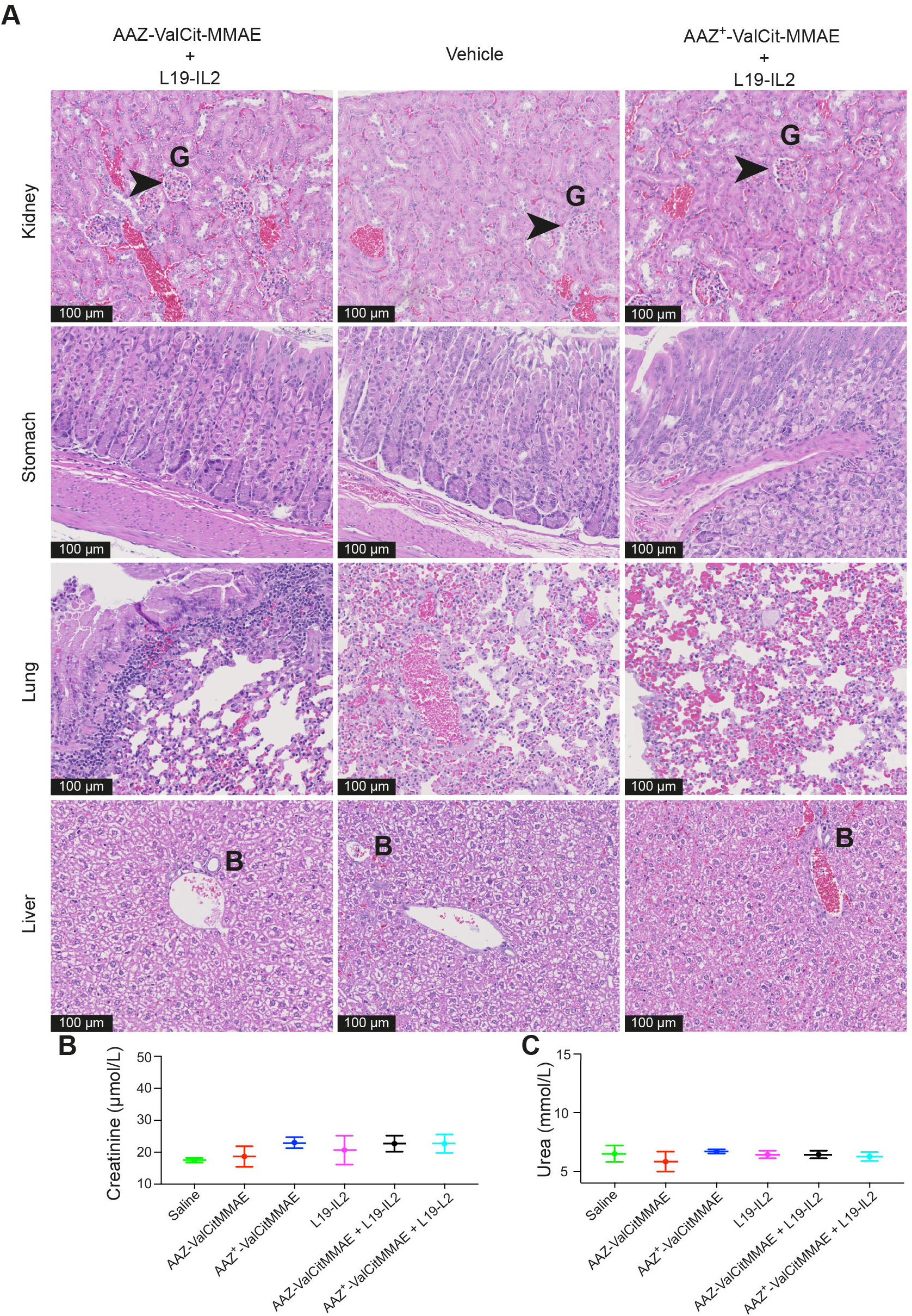
*Ex vivo* histological analysis and plasma analysis after therapy in BALB/c *nu/nu* mice treated with AAZ-ValCit-MMAE or AAZ^+^-ValCit-MMAE in combination with L19-IL2. (**A**) Representative pictures of H&E staining on kidney, stomach, lung and liver samples from different treated groups of mice collected three months after first exposure to the drugs (black scale bar = 100 μm, B = bile ducts in the liver, G = glomeruli structures in the kidney, arrow = inflammatory infiltrate with increase of fibrous tissue). (**B**) Creatinine and (**C**) urea plasma concentration measured three months after treatment with SMDC products, L19-IL2 and combinations of the drugs (n = 3 per group). Administration of SMDCs and immunocytokines did not interfere with normal kidney function as demonstrated by physiological levels of creatinine and urea in plasma.

### Analysis of chronic toxicity of combination treatment of SMDCs with L19-IL12

Gross examination of mice revealed a moderately enlarged spleen in all animals treated with L19-IL12 alone or in combination with AAZ^+^-ValCit-MMAE compared to the control group. In the liver, both treatment groups exhibited a mild [40] periportal infiltrate of mononuclear cells [**Figure 6**], rarely associated with increased connective tissue. Both changes were not noted in the control animal and did not differ significantly in between the groups.

**Figure 6:**
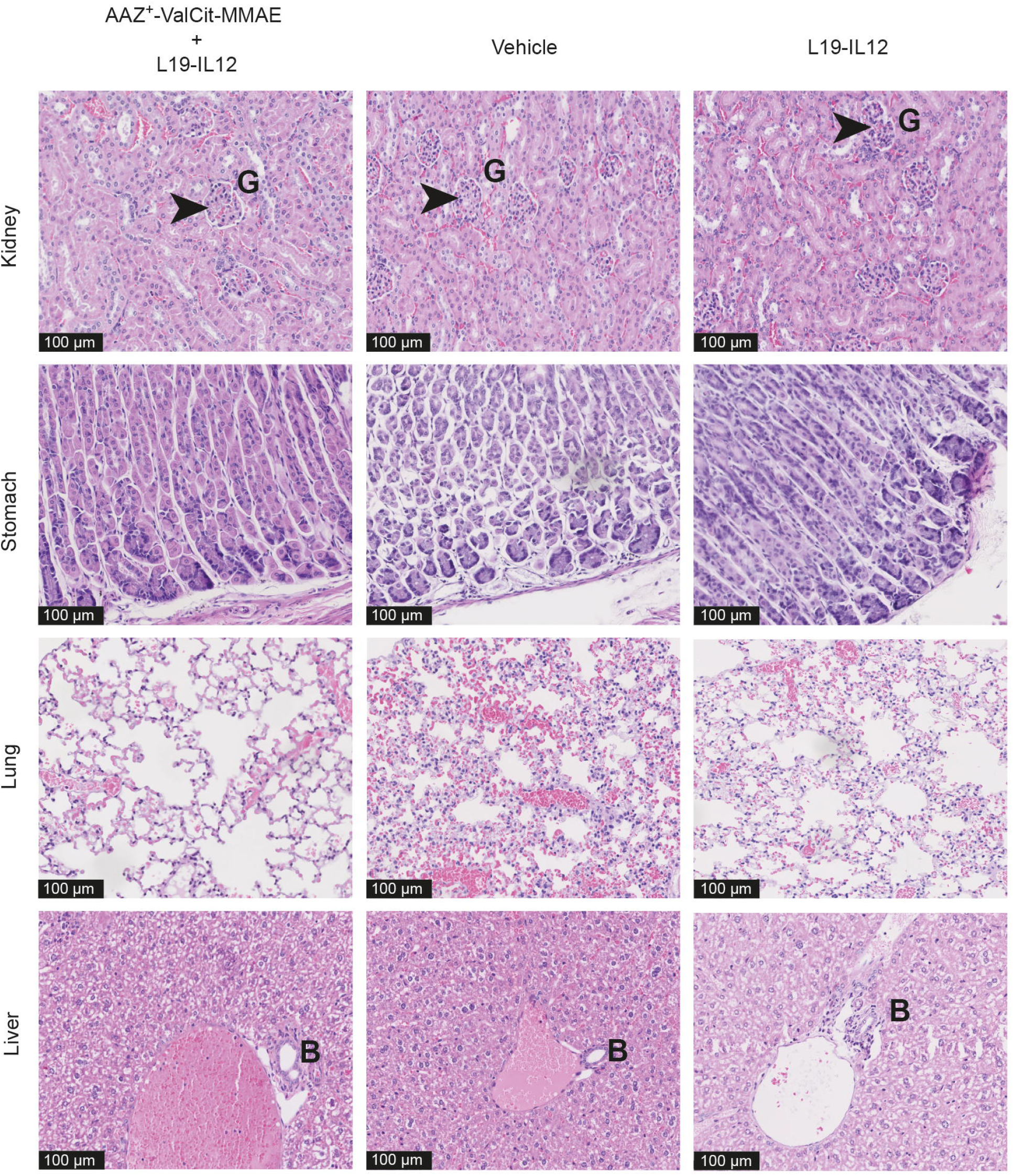
*Ex vivo* histological analysis after therapy in BALB/c *nu/nu* mice treated with AAZ^+^-ValCit-MMAE in combination with L19-IL12 or with L19-IL12 alone. Representative pictures of H&E staining on kidney, stomach, lung and liver samples from different treated groups of mice collected three months after first exposure to the drugs (black scale bar = 100 μm, B = bile ducts in the liver, G = glomeruli structures in the kidney, arrow = inflammatory infiltrate with increase of fibrous tissue).

The histopathological examination of kidney, lung and stomach did not show signs of inflammation and /or fibrosis [**Figure 6**]. In the spleen, a marked increase in extramedullary hematopoiesis in the red pulp correlated macroscopically with an enlarged spleen. The bone marrow of both treatment groups revealed a high cellularity with all three cell lines: erythropoiesis, myelopoiesis and thrombocytopoiesis were present with maturation in all three lines and did not seem to be altered compared to the control animal. [**Supplementary information**].

## Discussion

We have recently demonstrated that the anti-cancer activity of AAZ^+^-ValCit-MMAE, a small molecule-drug conjugate targeting carbonic anhydrase IX, is potently enhanced by the combination with targeted interleukin-2 [31]. In this article we have extended our previous findings, describing the *in vivo* anti-cancer activity and safety profile of novel therapies consisting of SMDC products targeting carbonic anhydrase IX combined with PD-1 blockade and with the antibody-cytokine fusions L19-IL2 and L19-IL12. Combination with all immunomodulatory drugs tested enhanced the therapeutic activity of small molecule-drug conjugates. The result of a systematic histopathological evaluation in treated mice shows that curative doses of SMDCs in combination with L19-IL2 and with L19-IL12 do not cause chronic toxicity in kidney and other healthy organs.

The clinical efficacy of targeted cytotoxics strictly depends on the amount of payload that can be delivered to the tumor cells, compatibly with a safe administration to the patients. While being designed to specifically deliver their toxic payload to tumors, both ADCs and SMDCs are found at significantly high concentrations in non-target organs [41]. Antibodydrug conjugates are slowly cleared by the liver. Common clinical side effects of ADCs include liver damage, bone marrow toxicity and hemorrhage [12]. Small molecule-drug conjugates are rapidly cleared by the kidney, with very short plasma half-lives and time of exposure in other healthy organs. The toxicity profile of SMDCs has not been sufficiently investigated, with only few preclinical [42,43] and clinical [44] examples available at present. We have demonstrated that anti-CAIX SMDC products are safe when administered at curative doses, with no signs of chronic damage in non-target healthy organs and in those healthy structures known to express CAIX [27],.These findings are in strong contrast with what has been reported for small molecule-radio conjugates, for which administration of therapeutic doses typically results in significant kidney toxicity [45]. Our results suggest that the delivery of cytotoxic drugs damaging cells in rapid proliferation may be advantageous over the use of therapeutic radionuclide payloads that damage indistinctively all type of cells [46].

Immunocytokines and checkpoint inhibitors have previously been found to potently enhance the anti-cancer activity of untargeted chemotherapy [39,47] and of antibody-drug conjugates [48]. When given as combination, the administration of immunotherapy and of cytotoxic drugs can be harmonized to maximize therapeutic potential and minimize systemic toxicities. we found that AAZ-ValCit-MMAE and AAZ^+^-ValCit-MMAE administered before immunotherapy were more efficacious than immunotherapy first followed by SMDCs, We suggest that the initial damage induced by targeted cytotoxics can render the tumor microenvironment more susceptible to the action of immunotherapy, as cytotoxic agents increase the expression of stress proteins such as Mic-A [39,49] and cause immunogenic cell death [50].

Our results provide a rationale for the clinical development of anti-CAIX small moleculedrug conjugates in combination with immunocytokines and PD-1 inhibitors. Small molecule-drug conjugates may represent a valuable alternative to antibody-drug conjugates, with clear advantages in terms of tumor targeting performance [25] and ease of manufacture [10].

## Acknowledgements

The authors wish to thank Baptiste Gouyou and Tiziano Ongaro for their technical support during *in vivo* experiments and Theresa Pesch for technical support with necropsy and histology. D.N. acknowledges funding from ETH Zurich. This project has received funding from the Swiss National Science Foundation (Grant Nr. 310030_182003/1) and the European Research Council (ERC) under the European Union’s Horizon 2020 research and innovation program (grant agreement 670603).

This project has received funding from the European Union’s Horizon 2020 research and innovation programme under the Marie Skłodowska-Curie grant agreement No 861316

