## Supplementary information for "Immunotherapy with immunocytokines and PD-1 blockade enhances the anticancer activity of small molecule-drug conjugates targeting carbonic anhydrase IX"

a) Philochem AG, Libernstrasse 3, CH-8112 Otelfingen (Switzerland)

b) Laboratory for Animal Model Pathology, Universität Zürich, Winterthurerstrasse 268, CH-8057 Zurich (Switzerland)

c) Department of Chemistry and Applied Biosciences, Swiss Federal Institute of Technology (ETH Zürich), Vladimir-Prelog-Weg 4, CH-8093 Zurich (Switzerland)

*) Corresponding author

Running Title:

Immunotherapy in combination with anti-CAIX SMDCs

Keywords:

Tumor Targeting, Small Molecule-Drug Conjugates, Immunotherapy, Carbonic Anhydrase IX, Therapy Studies

Financial Support:

D.N. acknowledges funding from ETH Zurich. This project has received funding from the Swiss National Science Foundation (Grant Nr. 310030_182003/1) and the European Research Council (ERC) under the European Union's Horizon 2020 research and innovation program (grant agreement 670603).

Disclosure of Potential Conflict of Interest:

D.N. is a co-founder and shareholder of Philogen (www.philogen.com), a Swiss-Italian Biotech company that operates in the field of ligand-based pharmacodelivery. J.M., S.D., A.V. and S.C. are employees of Philochem AG, daughter company of Philogen acting as discovery unit of the group.

Submitted as Research Article to *Molecular Cancer Therapeutics*

#### Abbreviations

| | |
|---|---|
| AA | Antibiotic Antimycotic |
| AAZ | Acetazolamide |
| $AAZ^+$ | Acetazolamide plus |
| Arg | Arginine |
| Asp | Aspartic Acid |
| Cit | Citrulline |
| CT | Chlorotrityl |
| Cys | Cysteine |
| CuAAC | Azide-Alkyne Huisgen Cycloaddition |
| DIPEA | *N*,*N*-Diisopropylethylamine |
| DMF | *N*,*N*-Dimethylformamide |
| DMSO | Dimethylsulfoxide |
| ESI | Electrospray Ionization |
| eq | Equivalents |
| FA | Formic Acid |
| FCS | Fetal Calf Serum |
| Fmoc | 9-Fluorenylmethoxycarbonyl |
| Glu | Glutamic acid |
| HATU | O-(7-azabenzotriazol-1-yl)-tetramethyl-uronium hexafluorophosphate |
| HBSS | Hank's Balanced Salt Solution |
| HOBt | Hydroxybenzotriazole |
| HPLC | High Performance Liquid Chromatography |
| LC/MS | Liquid Chromatography / Mass Spectrometry |
| Lys | Lysine |
| *m/z* | mass-to-charge ratio |

| | |
|---|---|
| MeCN | Acetonitrile |
| MMAE | Monomethyl Auristatin E |
| MS | Mass Spectroscopy |
| MW | Molecular Weight |
| PAB | *p*-aminobenzylalcohol |
| Pbf | 2,2,4,6,7-pentamethyldihydrobenzofuran-5-sulfonyl |
| PBS | Phosphate-Buffered Saline |
| ppm | Part per million |
| RCC | Renal Cell Carcinoma |
| RP | Reverse Phase |
| RPMI | Roswell Park Memorial Institute |
| r.t. | Room Temperature |
| SD | Standard Deviation |
| SEM | Standard Error of the Mean |
| SMDC | Small Molecule-Drug Conjugate |
| SPPS | Solid Phase Peptide Synthesis |
| *t*Bu | *tert*-Butyl |
| TBTA | Tris(benzyltriazolylmethyl)amine |
| TFA | Trifluoroacetic Acid |
| THF | Tetrahydrofuran |
| TIPS | Triisopropylsilane |
| Trt | Trityl |
| Val | Valine |

#### General Remarks and Procedures

Peptide grade *N,N*-dimethylformamide (DMF) for solid phase synthesis was bought from ABCR. All other solvents were used as supplied by Merck or Sigma Aldrich in HPLC or analytical grade. H-Cys(Trt)-2-CT-polystyrene resin was purchased from RAPP Polymere (Ernst-Simon-Strasse 9, 72072 Tuebingen, Germany). Maleimidocaproyl-ValCit-*p*-aminobenzylalcohol-MMAE was purchased from Levena Biopharma (No.9 Weidi Road, Qixia District, Nanjing, 210046, China). L19-IL2 was produced by Philogen S.p.A. (Via Bellaria, 35, 53018 Sovicille SI, Italy) and diluted to the concentration used for therapy studies with the appropriate formulation buffer (Philogen). Anti-mouse-PD1 antibody was purchased from Bio x Cell (10 Technology Dr., Suite 2B West Lebanon, NH 03784, US). All other reagents were purchased from Sigma Aldrich, Fluorochem, Chemimpex or TCI and used as supplied. Yields refer to chromatographically purified and spectroscopically pure compounds, unless noted otherwise.

Liquid-Chromatography/Mass-Spectrometry (LC/MS) spectra presented in the **Supplementary Information** were recorded on an Agilent 6100 Series Single Quadrupole MS system combined with an Agilent 1200 Series LC, using an InfinityLab Poroshell 120 EC-C18 Column, 2.7 µm, 4.6 × 50 mm at a flow rate of 0.6 ml min$^{-1}$, 10% MeCN in 0.1% aq. FA to 100% MeCN in 6 min.

Reversed-phase high-pressure liquid chromatography (RP-HPLC) were performed on a Agilent 1200 Series RP-HPLC with PDA UV detector, using a Synergi 4µm, Polar-RP 80Å 10 × 150 mm C18 column at a flow rate of 5 ml min$^{-1}$ with linear gradients of solvents A and B (A = Millipore water with 0.1% trifluoroacetic acid [TFA], B = MeCN with 0.1% trifluoroacetic acid [TFA]).

Size-exclusion chromatography was performed on a Superdex 200 Increase 10/300 GL column on an ÄKTA FPLC (GE Healthcare).

#### Synthesis of Compound 1

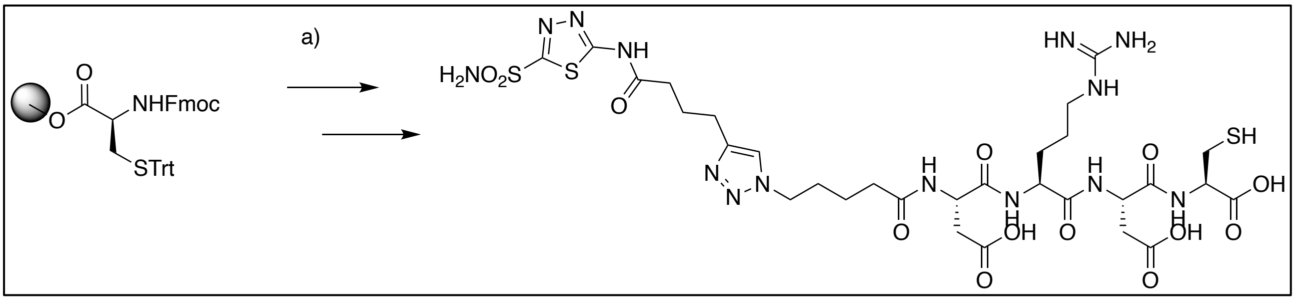

*a)* Fmoc-Glu-(O*t*Bu)-OH , HOBt, HBTU, DIPEA, DMF, 1h, r.t. ; Fmoc-Arg-(Pbf)-OH , HOBt, HBTU, DIPEA, DMF, 1h, r.t. ; Fmoc-Glu-(O*t*Bu)-OH , HOBt, HBTU, DIPEA, DMF, 1h, r.t. ; 5-azido-pentanoic acid , HOBt, HBTU, DIPEA, DMF, 1h, r.t. ; 4,4-bis(4-hydroxyphenyl)valeric acid, HOBt, HBTU, DIPEA, DMF, 1h, r.t. ; *N*-(5-sulfamoyl-1,3,4-thia-diazol-2-yl)hex-5-ynamide, CuI, TBTA, DMF/THF 1:1, overnight, r.t. *b)* TFA (74.5 %), m-cresol (3.5 %), thioanisol (3.5 %), TIPS (14%), water (3.5 %), 1h, r.t.

Commercially available H-Cys(Trt)-2-Chlorotritil resin (500 mg) was used for the synthesis of the peptide following the Fmoc-protocol. The peptide was cleaved from the resin with a cleavage cocktail composed by TFA (75.5%), m-Cresol (3.5%), Thioanisole (3.5%), Triisopropylsilane (14%) and mQ water (3.5%). Compound **1** was purified from the crude via reverse-phase HPLC. Product containing fractions were identified by mass spectrometry and lyophilized overnight to obtain compound **1** as a white powder (120 mg; 0.132 mmol; 35% yield).

<u>**Synthesis of AAZ-ValCit-MMAE**</u>

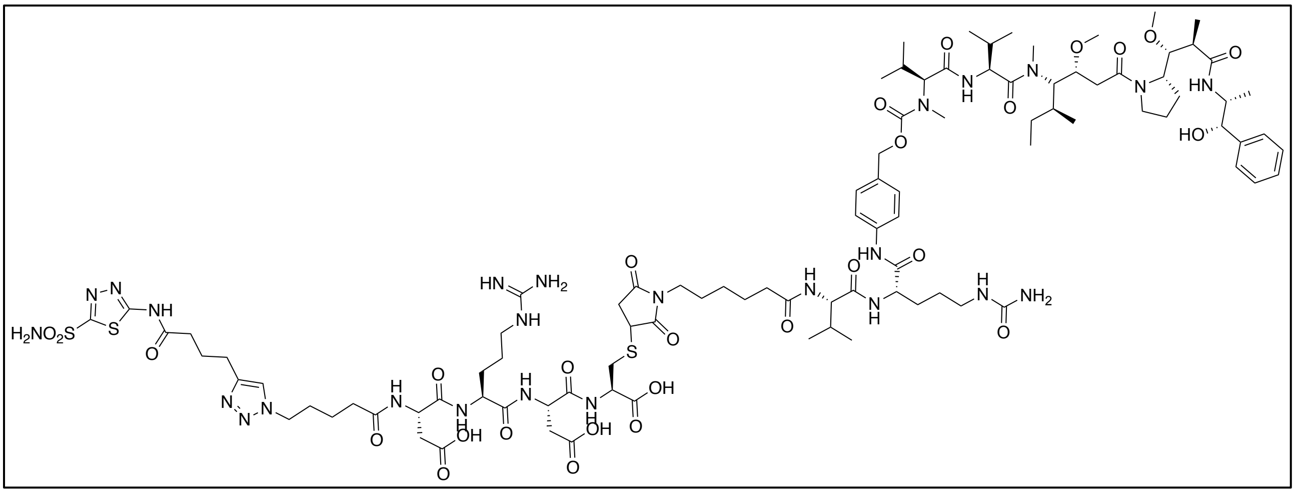

Compound **1** (3.4 mg; 3.7 µmol; 2.0 eq) was dissolved in degassed PBS (pH 7.4; 600 µl). MC-ValCit-MMAE (2.5 mg; 1.8 µmol; 1.0 eq) was added as a DMF solution (350 µl) and the mixture was stirred at room temperature. Product formation was monitored by LC/MS. MC-ValCit-MMAE was fully conjugated after 3 hours and the solvents were removed under vacuum. The crude was diluted in a 1:1 $H_2O$/MeCN mixture (1 ml) and purified by RP-HPLC. Product containing fractions were identified by mass spectrometry and lyophilized overnight to obtain **AAZ-ValCit-MMAE** as a white powder (1.18 mg; 0.59 µmol; 30% yield).

#### Synthesis of Compound 2

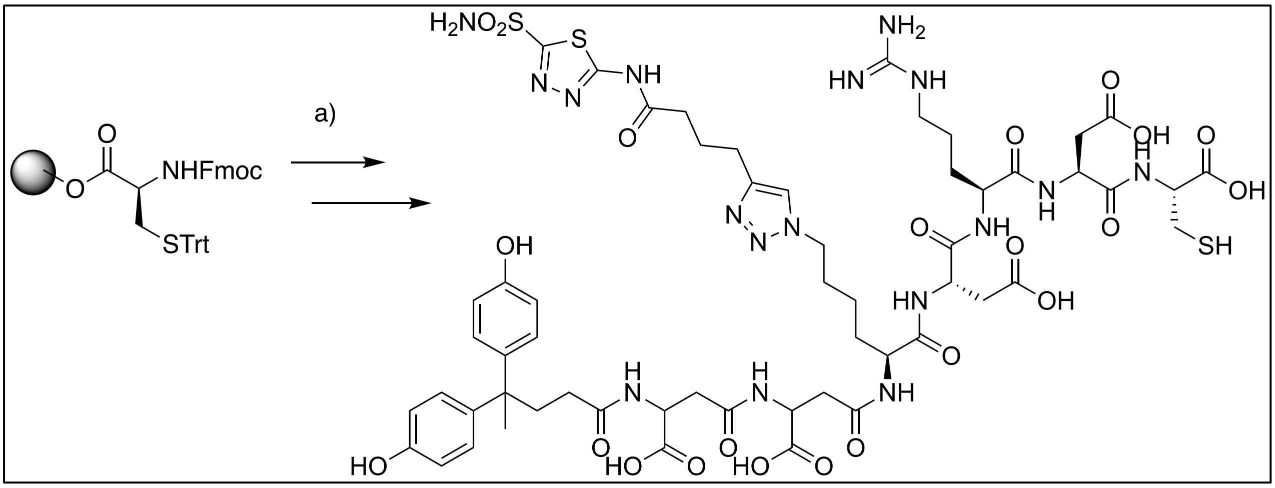

*a)* Fmoc-Glu-(O*t*Bu)-OH , HOBt, HBTU, DIPEA, DMF, 1h, r.t. ; Fmoc-Arg-(Pbf)-OH , HOBt, HBTU, DIPEA, DMF, 1h, r.t. ; Fmoc-Glu-(O*t*Bu)-OH , HOBt, HBTU, DIPEA, DMF, 1h, r.t. ; Fmoc-Lys-N3 , HOBt, HBTU, DIPEA, DMF, 1h, r.t. ; Fmoc-Glu-(αO*t*Bu)-OH , HOBt, HBTU, DIPEA, DMF, 1h, r.t. ; Fmoc-Glu-(αO*t*Bu)-OH , HOBt, HBTU, DIPEA, DMF, 1h, r.t. ; 4,4-bis(4-hydroxyphenyl)valeric acid, HOBt, HBTU, DIPEA, DMF, 1h, r.t. ; *N*-(5-sulfamoyl-1,3,4-thia-diazol-2-yl)hex-5-ynamide, CuI, TBTA, DMF/THF 1:1, overnight, r.t. *b)* TFA (74.5 %), m-cresol (3.5 %), thioanisol (3.5 %), TIPS (14%), water (3.5 %), 1h, r.t.

Commercially available H-Cys(Trt)-2-Chlorotritil resin (500 mg) was used for the synthesis of the peptide following the Fmoc-protocol. The peptide was cleaved from the resin with a cleavage cocktail composed by TFA (75.5%), m-Cresol (3.5%), Thioanisole (3.5%), Triisopropylsilane (14%) and mQ water (3.5%). Compound **1** was purified from the crude via reverse-phase HPLC. Product containing fractions were identified by mass spectrometry and lyophilized overnight to obtain compound **2** as a white powder (12 mg; 8.37 μmol; 3.5% yield).

#### Synthesis of AAZ+-ValCit-MMAE

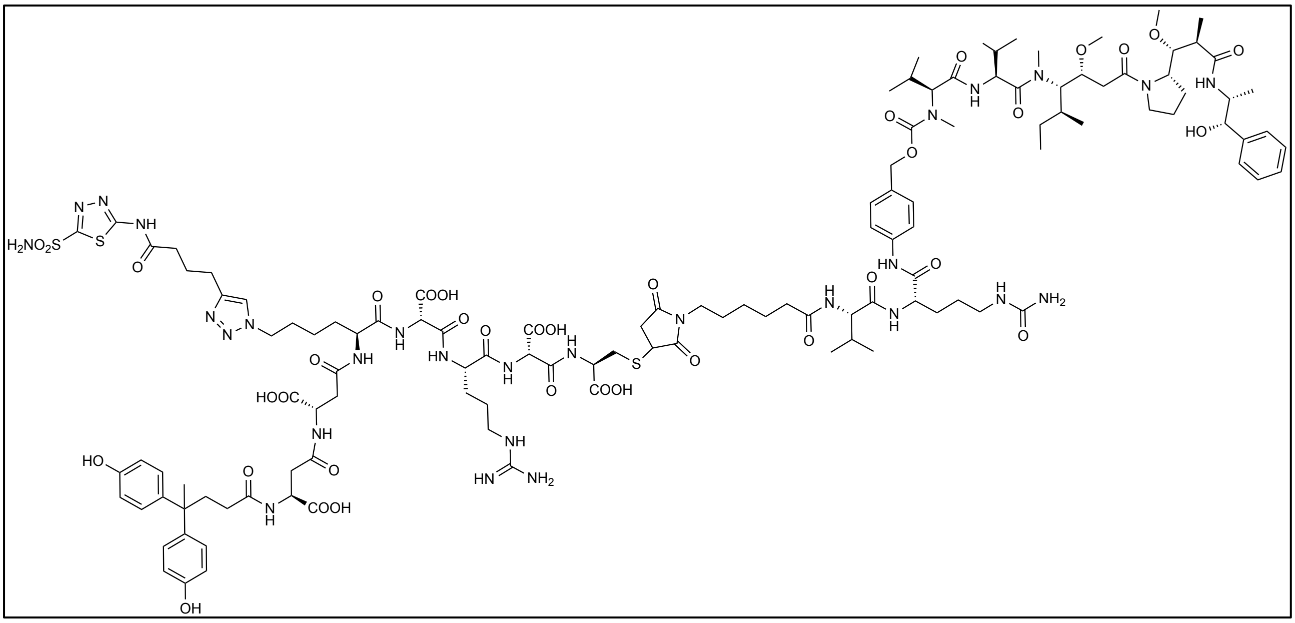

Compound **1** (6.0 mg; 6.6 µmol; 4.0 eq) was dissolved in degassed PBS (pH 7.4; 600 µl). MC-ValCit-MMAE (2.2 mg; 1.6 µmol; 1.0 eq) was added as a DMF solution (350 µl) and the mixture was stirred at room temperature. Product formation was monitored by LC/MS. MC-ValCit-MMAE was fully conjugated after 3 hours and the solvents were removed under vacuum. The crude was diluted in a 1:1 $H_2O$/MeCN mixture (1 ml) and purified by RP-HPLC. Product containing fractions were identified by mass spectrometry and lyophilized overnight to obtain **AAZ+-ValCit-MMAE** as a white powder (1.26 mg; 0.59 µmol; 53% yield).

### HPLC profiles

*Compound **1***

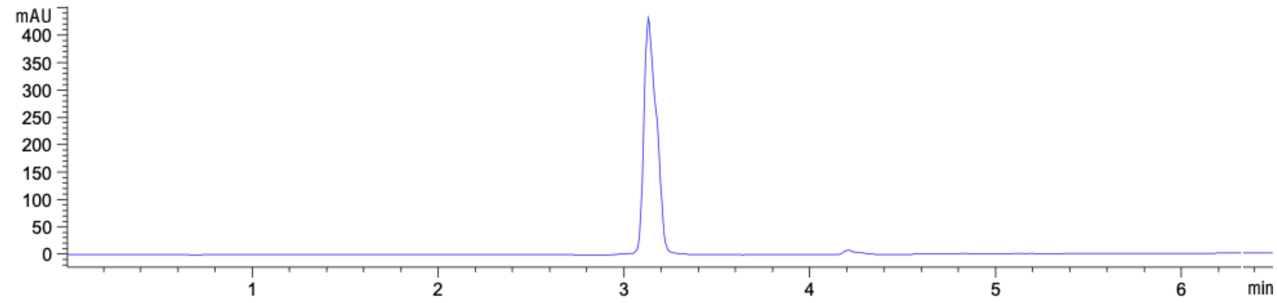

MS (ESI) *m/z* calcd. for $[C_{30}H_{46}N_{14}O_{13}S_3]^+$: 907.25 $[M+H]^+$, found: 907.20.

*AAZ-ValCit-MMAE*

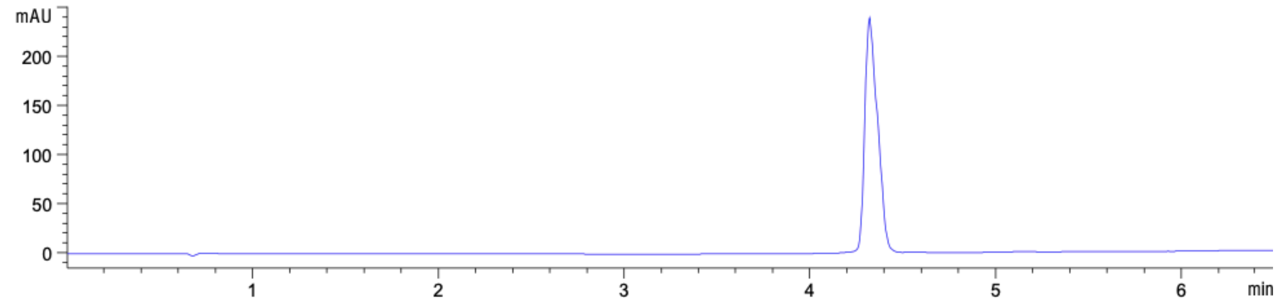

MS (ESI) *m/z* calcd. for $[C_{98}H_{151}N_{25}O_{28}S_3]^+$: 2223.02 $[M+2H]^{2+}$, found: 2223.00.

*Compound **2***

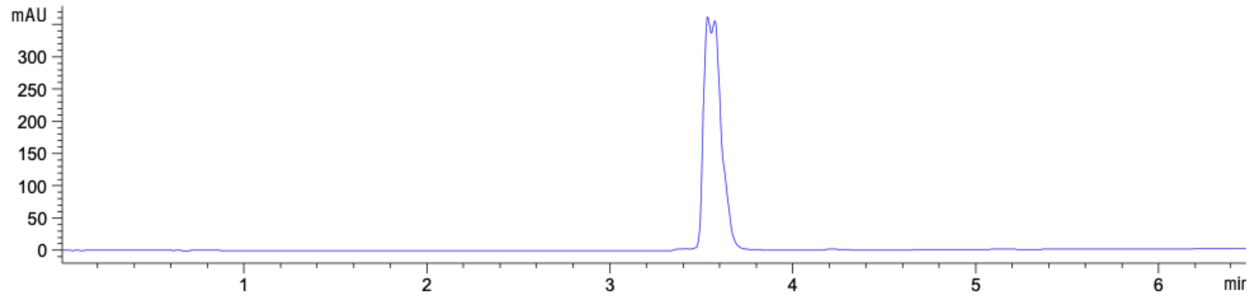

MS (ESI) *m/z* calcd. for $[C_{56}H_{76}N_{17}O_{22}S_3]^+$: 1433.44 $[M+H]^+$, found: 1433.00.

*AAZ⁺-ValCit-MMAE*

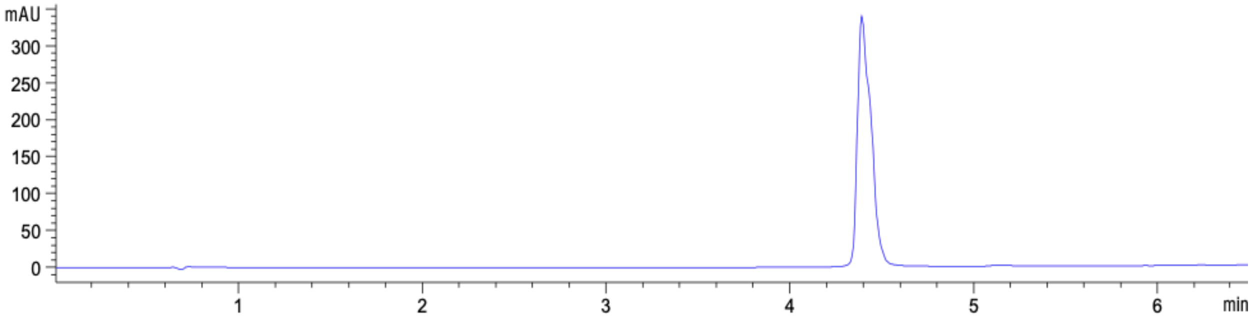

MS (ESI) *m/z* calcd. for $[C_{124}H_{182}N_{28}O_{37}S_3]^{2+}$: 1375.17 $[M+2H]^{2+}$, found: 1375.00.

**L19-IL2 quality control**

Size-exclusion chromatography and LC/MS analysis were performed in order to confirm the quality of the protein.

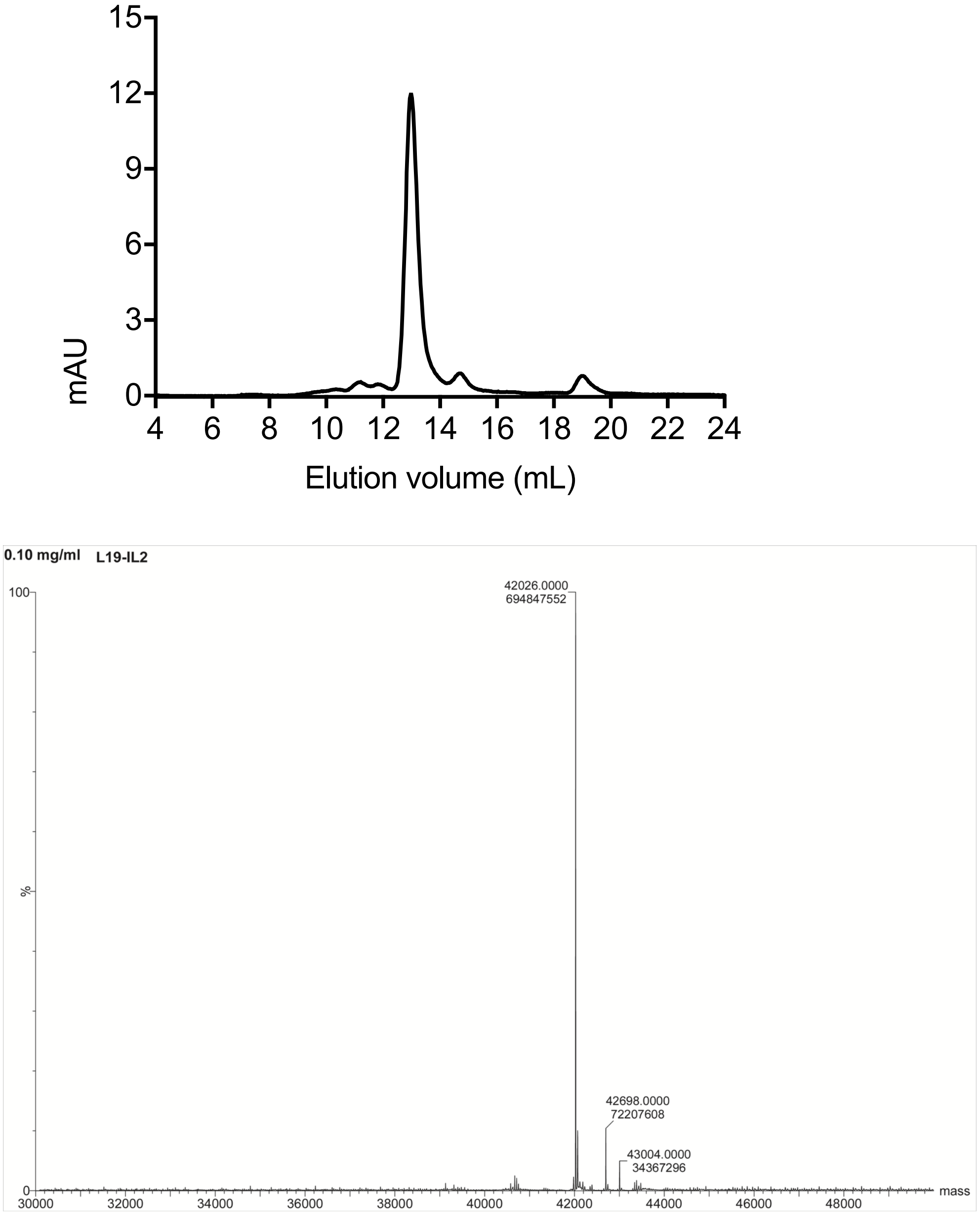

**Supplementary Figure 1:** Quality control of L19-IL2: size-exclusion chromatography and LC/MS profiles.

**L19-IL12 sequence**

**NheI**// Leader Sequence-mIL12 beta p40-Linker-mIL12 alpha p35- Linker- L19 -Linker –L19 //**NotI**

CTCCGCTAGCGTCGACCATGGGCTGGAGCCTGATCCTCCTGTTCCTCGTCGCTGTGGC-

TACAGGTGTGCACTCGATGTGGGAGCTGGAGAAAGACGTTTATGTTGTAGAGGTGGAC

TGGACTCCCGATGCCCCTGGAGAAACAGTGAACCTCACCTGTGACACGCCTGAAGAA-

GATGACATCACCTGGACCTCAGACCAGAGACATGGAGTCATAGGCTCTGGAAAGACCC

TGACCATCACTGTCAAAGAGTTTCTAGATGCTGGCCAGTACAC-

CTGCCACAAAGGAGGCGA-

GACTCTGAGCCACTCACATCTGCTGCTCCACAAGAAGGAAAATGGAATTTGGTCCACT

GAAATTTTAAAAAATTTCAAAAACAAGACTTTCCTGAAGTGTGAAGCAC-

CAAATTACTCCG-

GACGGTTCACGTGCTCATGGCTGGTGCAAAGAAACATGGACTTGAAGTTCAACATCAA

GAGCAGTAGCAGTTCCCCTGACTCTCGGGCAGTGACATGTG-

GAATGGCGTCTCTGTCTG-

CAGAGAAGGTCACACTGGACCAAAGGGACTATGAGAAGTATTCAGTGTCCTGCCAGGA

GGATGTCACCTGCCCAACTGCCGAGGAGACCCTGCCCATTGAACTGGCGTTGGAA-

GCACGG-

CAGCAGAATAAATATGAGAACTACAGCACCAGCTTCTTCATCAGGGACATCATCAAAC

CAGACCCGCCCAAGAACTTGCAGATGAGGCCTTTGAAGAACTCACAGGTG-

GAGGTCAGCTGG-

GAGTACCCTGACTCCTGGAGCACTCCCCATTCCTACTTCTCCCTCAAGTTCTTTGTTCGA

ATCCAGCGCAAGAAAGAAAAGATGAAGGAGACAGAGGAGGGGTGTAAC-

CAGAAAGGTGCGTTCCTCGTAGAGAGGACATCTACCGAAGTCCAATGCAAAGGCGGG

AATGTCTGCGTGCAAGCTCAGGATCGCTATTACAATTCCTCGTGCAGCAAGTGGG-

CATGTGTTCCCTGCAGGGTCCGATCCGGTGGAGGCGGTTCAGGCGGAGGTGGCTCTGG

CGGTGGCGGATCGAGGGTCATTCCAGTCTCTGGACCTGCCAGGTGTCTTAGCCAG-

TCCCGAAACCTGCTGAAGACCACAGATGACATGGTGAAGACGGCCAGAGAA***AAGCTT***A

AACATTATTCCTGCACTGCTGAAGACATCGATCATGAAGACATCACACGGGACCAAAC-

CAG-

CACATTGAAGACCTGTTTACCACTGGAACTACACAAGAACGAGAGTTGCCTGGCTACT

AGAGAGACTTCTTCCACAACAAGAGGGAGCTGCCTGCCCCCACAGAAGACGTCTTT-

GATGATGACCCTGTGCCTTGGTAGCATCTATGAGGACTTGAAGATGTACCAGACAGAG

TTCCAGGCCATCAACGCAGCACTTCAGAATCACAACCATCAGCAGATCATTCTAGA-

CAAGGG-

CATGCTGGTGGCCATCGATGAGCTGATGCAGTCTCTGAATCATAATGGCGAGACTCTG

CGCCAGAAACCTCCTGTGGGAGAAGCAGACCCTTACAGAGTGAAAATGAAGCTCTG-

CATCCTGCTTCACGCCTTCAGCACCCGCGTCGTGACCATCAACAGGGTGATGGGCTATC

TGAGCTCCGCCGGTAGCGCTGATGGAGAGGTGCAGCTGTTGGAGTCTGGGG-

GAGGCTTGG-

TACAGCCTGGGGGGTCCCTGAGACTCTCCTGTGCAGCCTCTGGATTCACCTTTAGCAGT

TTTTCGATGAGCTGGGTCCGCCAGGCTCCAGGGAAGGGGCTGGAG-

TGGGTCTCATCTATTAG-

TGGTAGTTCGGGTACCACATACTACGCAGACTCCGTGAAGGGCCGGTTCACCATCTCC

AGAGACAATTCCAAGAACACGCTGTATCTGCAAATGAACAGCCTGAGAGCCGAG-

GACAC-

GGCCGTATATTACTGTGCGAAACCGTTTCCGTATTTTGACTACTGGGGCCAGGGAACCC

TGGTCACCGTCTCGAGTGGGTCCAGTGGCGGTGAAATTGTGTTGACGCAG-

TCTCCAGGCACCCTGTCTTTGTCTCCAGGGGAAAGAGCCACCCTCTCCTGCAGGGCCAG

TCAGAGTGTTAGCAGCAGCTTTTTAGCCTGGTACCAGCAGAAAC-
CTGGCCAGGCTCCCAGGCTCCTCATCTATTATGCATCCAGCAGGGCCACTGGCATCCCA
GACAGGTTCAGTGGCAGTGGGTCTGGGACAGACTTCACTCTCACCATCAGCAGACTG-
GAGCCTGAAGATTTTGCAGTGTATTACTGTCAGCAGACGGGTCGTATTCCGCCGACGTT
CGGC*CAAGGGACCAAGGTGGAAATCAAA*TCTTCCTCATCCGGAAGTAGCTCTTCG***G-***
***GATCC***TCGTCCAGCGGCGAGGTGCAGCTGTTGGAGTCTGGGGGAGGCTTGGTACAGCC
TGGGGGGTCCCTGAGACTCTCCTGTGCAGCCTCTGGATTCACCTTTAGCAG-
TTTTTCGATGAGCTGGGTCCGCCAGGCTCCAGGGAAGGGGCTGGAGTGGGTCTCATCT
ATTAGTGGTAGTTCGGGTACCACATACTACGCAGACTCCGTGAAGGGCCGGTTCAC-
CATCTCCAGAGACAATTCCAAGAACACGCTGTATCTGCAAATGAACAGCCTGAGAGCC
GAGGACACGGCCGTATATTACTGTGCGAAACCGTTTCCGTATTTTGAC-
TACTGGGGCCAGGGAACCCTGGTCACCGTCTCGAGTGGGTCCAGTGGCGGTGAAATTG
TGTTGACGCAGTCTCCAGGCACCCTGTCTTTGTCTCCAGGGGAAA-
GAGCCACCCTCTCCTG-
CAGGGCCAGTCAGAGTGTTAGCAGCAGCTTTTTAGCCTGGTACCAGCAGAAACCTGGC
CAGGCTCCCAGGCTCCTCATCTATTATGCATCCAGCAGGGCCACTGGCATCCCAGA-
CAGGTTCAGTGGCAGTGGGTCTGGGACAGACTTCACTCTCACCATCAGCAGACTGGAG
CCTGAAGATTTTGCAGTGTATTACTGTCAGCAGACGGGTCGTATTCCGCCGAC-
GTTCGGC*CAAGGGACCAAGGTGGAAATCAAA*TAGTGA**GCGGCCGC**AAAAGGAAAA

#### L19-IL12 quality control

SDS-page and size-exclusion chromatography were performed in order to confirm the quality of the protein.

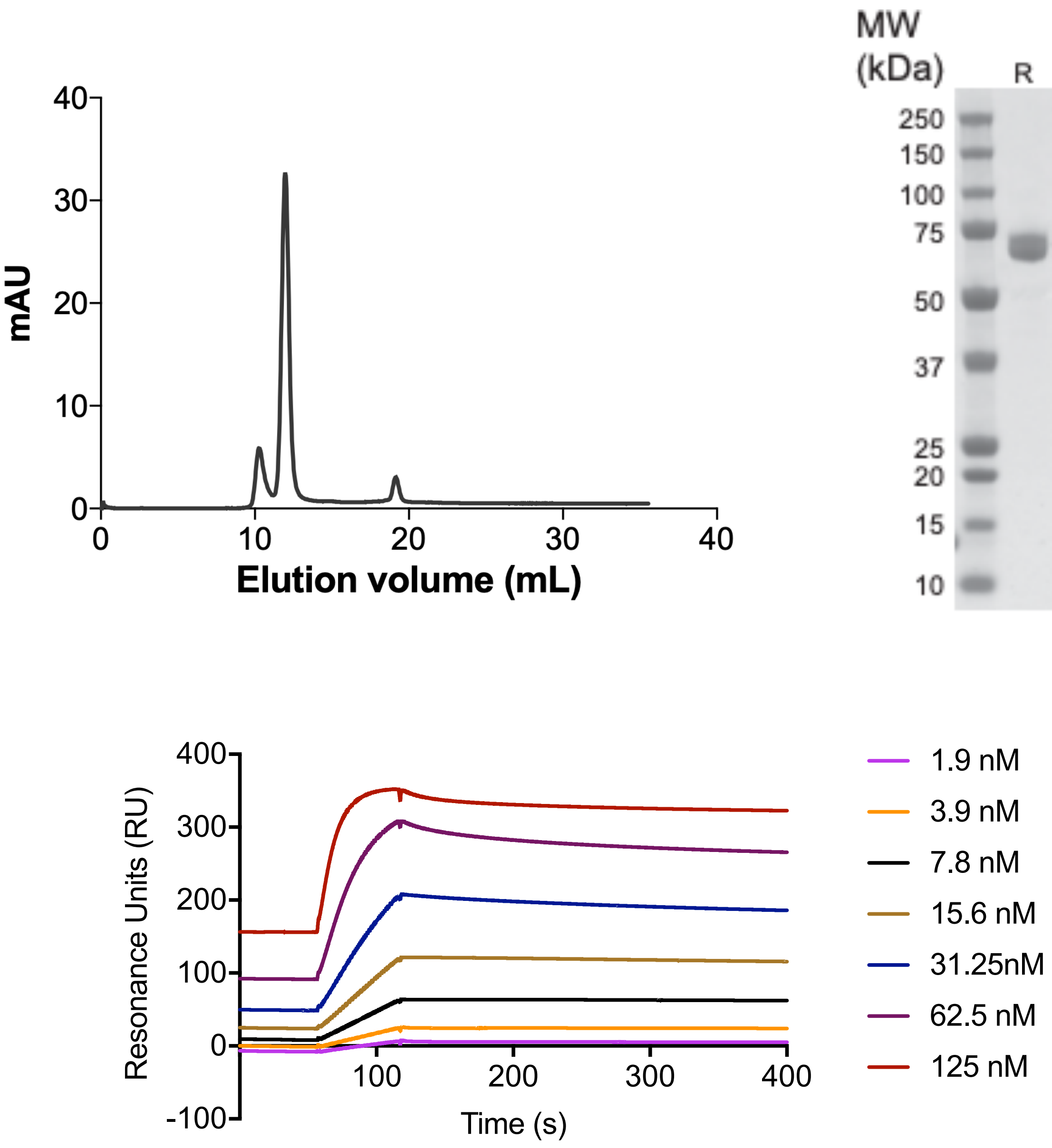

**Supplementary Figure 2:** Quality control of L19-IL12: size-exclusion chromatography profile, SDS-PAGE and binding to the cognate antigen EDB by Surface Plasmon Resonance.

#### Dose escalation of L19-IL12 in the SKRC-52 model

A dose escalation study was performed in athymic BALB/c *nu/nu* mice (8-10 weeks of age, Janvier) injecting four different doses (0.6, 0.8, 1.0 and 1.2 mg/kg) of the immunocytokine L19-IL12 at the day 1, 3 and 5. Toxicity was assessed by daily monitoring the body weight of the mice for 6 consecutive days. No acute toxicity was observed for L19-IL12 in this setting for the tested doses.

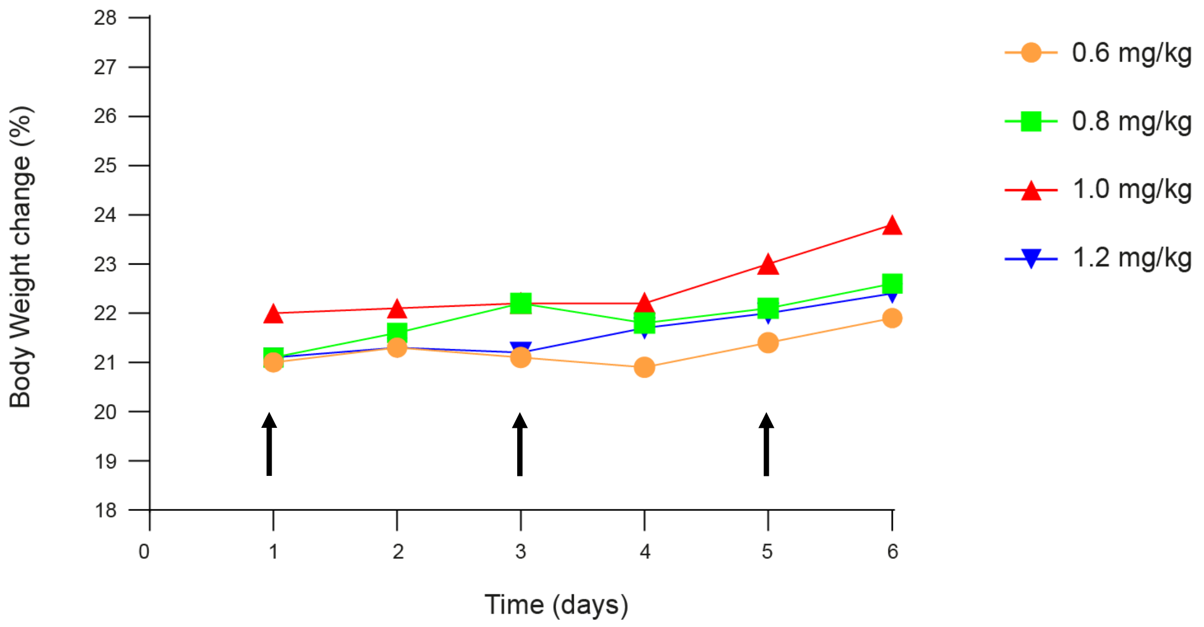

**Supplementary Figure 3:** Dose escalation of L19-IL12 in athymic BALB/c *nu/nu* mice (8-10 weeks of age, Janvier)

#### Statistical analysis of therapy experiments

Differences in tumor volume between therapeutic groups were compared using the two-way ANOVA analysis with Bonferroni post-test of Graphpad Prism 6 (La Jolla, CA, USA).

#### Therapy experiment: AAZ-ValCit-MMAE and AAZ$^+$-ValCit-MMAE combined with L19-IL2 with two different schedules

**Tumor Size (mm$^3$)**

*AAZ-ValCit-MMAE + L19-IL2* vs *AAZ$^+$-ValCit-MMAE + L19-IL2:*

no statistically significant difference

*AAZ-ValCit-MMAE + L19-IL2 vs. AAZ-ValCit-MMAE* → *L19-IL2:*

no statistically significant difference

*AAZ-ValCit-MMAE + L19-IL2 vs. AAZ$^+$-ValCit-MMAE,* → *L19-IL2:*

no statistically significant difference

*AAZ-ValCit-MMAE + L19-IL2 vs. vehicle:*

| | |
|---|---|
| Day 11 | $p < 0.01$ |
| From day 12 | $p < 0.0001$ |

*AAZ$^+$-ValCit-MMAE + L19-IL2* vs *AAZ-ValCit-MMAE* → L19-IL2:

no statistically significant difference

*AAZ$^{+}$-ValCit-MMAE + L19-IL2* vs *AAZ$^{+}$-ValCit-MMAE → L19-IL2*

no statistically significant difference

*AAZ$^{+}$-ValCit-MMAE + L19-IL2 vs. vehicle:*

| | |
|---|---|
| Day 10 | $p < 0.05$ |
| Day 11 | $p < 0.001$ |
| From day 12 | $p < 0.0001$ |

*AAZ-ValCit-MMAE → L19-IL2 vs. AAZ$^{+}$-ValCit-MMAE → L19-IL2:*

no statistically significant difference

*AAZ-ValCit-MMAE → L19-IL2 vs. vehicle:*

| | |
|---|---|
| Day 10 | $p < 0.05$ |
| Day 11 | $p < 0.001$ |
| From day 12 | $p < 0.0001$ |

*AAZ$^{+}$-ValCit-MMAE → L19-IL2 vs. vehicle:*

| | |
|---|---|
| Day 10 | $p < 0.05$ |
| Day 11 | $p < 0.001$ |
| From day 12 | $p < 0.0001$ |

**Therapy experiment: AAZ-ValCit-MMAE and AAZ$^{+}$-ValCit-MMAE combined with anti-PD-1**

**Tumor Size (mm$^3$)**

*AAZ-ValCit-MMAE vs. AAZ$^{+}$-ValCit-MMAE:*

no statistically significant difference

*AAZ-ValCit-MMAE vs.anti-PD-1:*

no statistically significant difference

*AAZ-ValCit-MMAE vs AAZ-ValCit-MMAE + anti-PD-1:*

| | |
|---|---|
| Day 22 | $p < 0.05$ |
| Day 24 | $p < 0.0001$ |

*AAZ-ValCit-MMAE vs. AAZ$^{+}$-ValCit-MMAE + anti-PD-1:*

| | |
|---|---|
| Day 24 | $p < 0.01$ |

*AAZ-ValCit-MMAE vs. Vehicle:*

no statistically significant difference

*AAZ$^{+}$-ValCit-MMAE vs.anti-PD1:*

no statistically significant difference

*AAZ$^+$-ValCit-MMAE vs. AAZ-ValCit-MMAE + anti-PD-1:*

| | |
|---|---|
| Day 24 | $p < 0.05$ |

*AAZ$^+$-ValCit-MMAE vs. AAZ$^+$-ValCit-MMAE + anti-PD-1:*

no statistically significant difference

*AAZ$^+$-ValCit-MMAE vs.Vehicle:*

no statistically significant difference

*Anti-PD-1 vs. AAZ-ValCit-MMAE + Anti-PD-1:*

| | |
|---|---|
| Day 22 | $p < 0.05$ |
| Day 24 | $p < 0.01$ |

*Anti-PD1 vs. AAZ$^+$-ValCit-MMAE + Anti-PD-1:*

| | |
|---|---|
| Day 24 | $p < 0.05$ |

*Anti-PD-1 vs. Vehicle:*

no statistically significant difference

*AAZ-ValCit-MMAE + anti-PD-1 vs. AAZ$^+$-ValCit-MMAE + anti-PD-1:*

no statistically significant difference

*AAZ-ValCit-MMAE + anti-PD-1 vs. Vehicle:*

| | |
|---|---|
| Day 22 | $p < 0.05$ |

Day 24 $p < 0.005$

*AAZ⁺-ValCit-MMAE + anti-PD-1 vs. Vehicle:*

Day 22 $p < 0.05$

Day 24 $p < 0.005$

**Therapy experiment: AAZ⁺-ValCit-MMAE combined with L19-IL12 (initial tumor size 100mm³)**

**Tumor Size ($mm^3$)**

*L19-IL12 vs. Vehicle:*

Day 14 $p < 0.05$

From day 15 $p < 0.0001$

*AAZ⁺-ValCit-MMAE vs. Vehicle:*

Day 15 $p < 0.05$

From day 16 $p < 0.01$

*AAZ⁺-ValCit-MMAE + L19-IL12 vs. Vehicle:*

Day 14 $p < 0.01$

From day 15 $p < 0.0001$

*AAZ⁺-ValCit-MMAE vs. L19-IL12:*

| | |
|---|---|
| Day 15 | $p < 0.05$ |
| Days 16, 17 and 18 | $p < 0.01$ |
| From day 19 | $p < 0.0001$ |

*$AAZ^+$-ValCit-MMAE vs $AAZ^+$-ValCit-MMAE → L19-IL12:*

| | |
|---|---|
| Day 14 | $p < 0.05$ |
| Days 15, 16, 17 and 18 | $p < 0.01$ |
| From day 19 | $p < 0.0001$ |

*L19-IL12 vs $AAZ^+$-ValCit-MMAE → L19-IL12:*

| | |
|---|---|
| Day 66 | $p < 0.01$ |

**Therapy experiment: $AAZ^+$-ValCit-MMAE combined with L19-IL12 (initial tumor size 200mm$^3$)**

#### Tumor Size (mm$^3$)

*L19-IL12 vs. Vehicle:*

no statistically significant difference

*$AAZ^+$-ValCit-MMAE vs. Vehicle:*

| | |
|---|---|
| From day 28 | $p < 0.01$ |

*$AAZ^+$-ValCit-MMAE → L19-IL12 vs. Vehicle:*

| | |
|---|---|
| Day 25 | $p < 0.01$ |
| Days 26 and 27 | $p < 0.001$ |
| From day 28 | $p < 0.0001$ |

*$AAZ^+$-ValCit-MMAE vs. L19-IL12:*

no statistically significant difference

*$AAZ^+$-ValCit-MMAE vs $AAZ^+$-ValCit-MMAE → L19-IL12:*

| | |
|---|---|
| Day 25 | $p < 0.05$ |
| Day 26 | no statistically significant difference |
| Day 27 | $p < 0.05$ |
| From day 30 | $p < 0.0001$ |

*L19-IL12 vs $AAZ^+$-ValCit-MMAE → L19-IL12:*

| | |
|---|---|
| Day 24 | $p < 0.05$ |
| Day 25 | $p < 0.01$ |
| Days 26 and 27 | $p < 0.005$ |
| From day 30 | $p < 0.0001$ |

#### Evaluation of chronic toxicity by *ex vivo* histology

SKRC-52 bearing mice treated with AAZ$^{+}$-ValCit-MMAE in combination with L19-IL12 or with L19-IL12 only (three per group) were sacrificed 90 days after treatment and fixed in formalin. Relevant organs were isolated and the microscopic morphology was analyzed compared to a corresponding control samples collected from a non-treated animal.

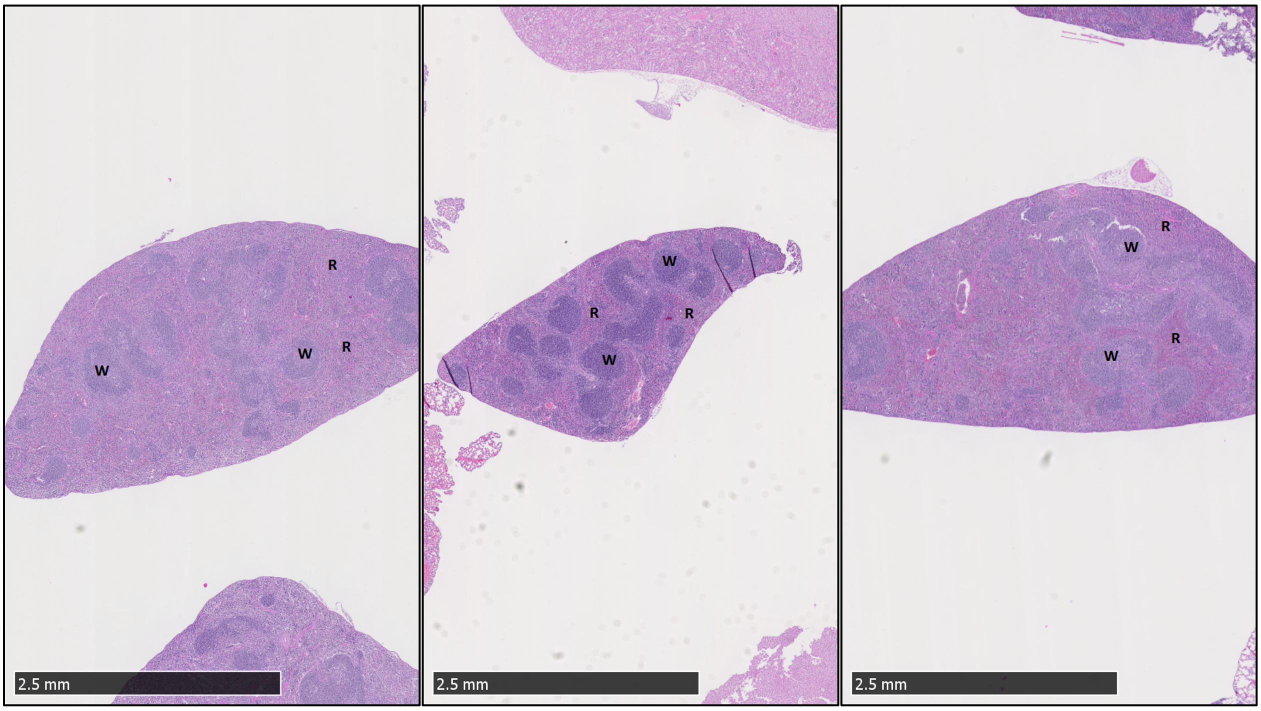

**Supplementary Figure 4**: Overview of cross section through the spleen demonstrating different size in control (center) and animals treated with L19-IL12 + AAZ$^{+}$-ValCit-MMAE (on the left) or with L19-IL12 (on the right). R= Red pulp, W= white pulp.

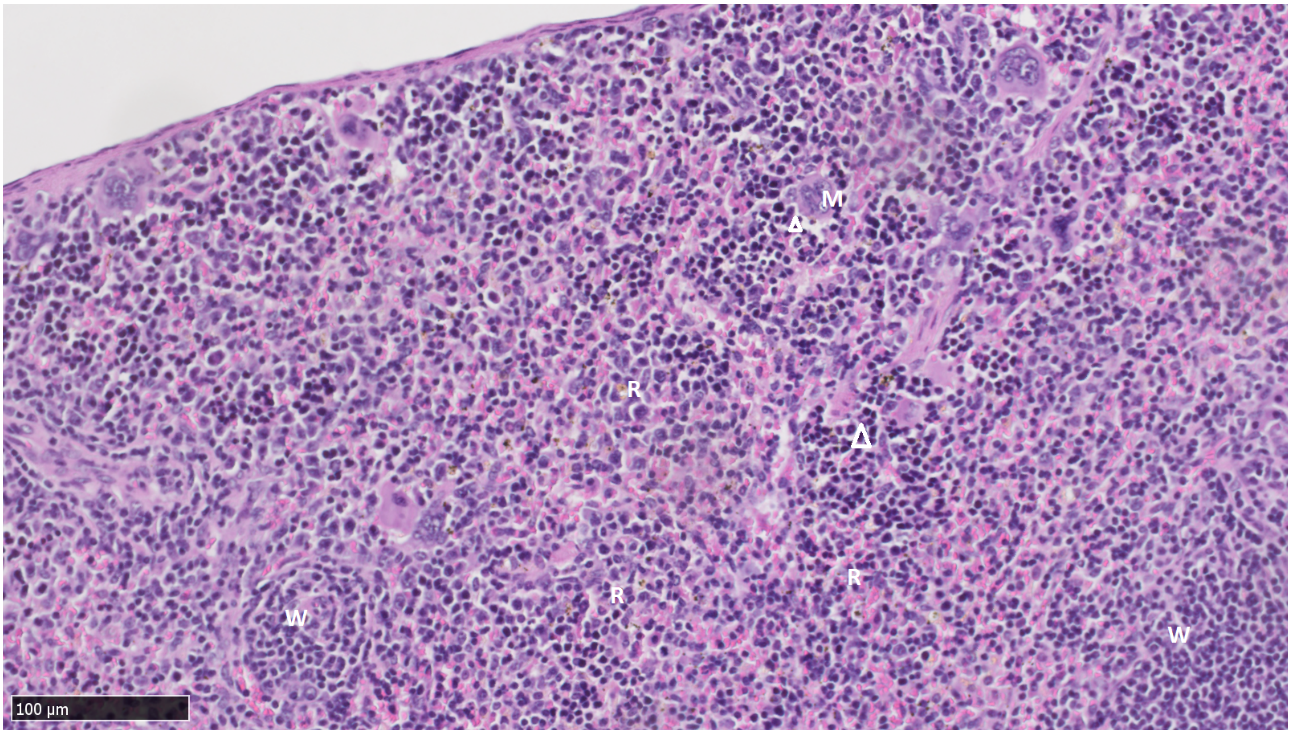

**Supplementary Figure 5:** Spleen close up analysis of mice treated with the combination L19-IL12 + AAZ+-ValCit-MMAE. W = white pulp, R = red pulp, M = Megakaryocytes and the arrow indicate extramedullary erythroid precursors in red pulp.

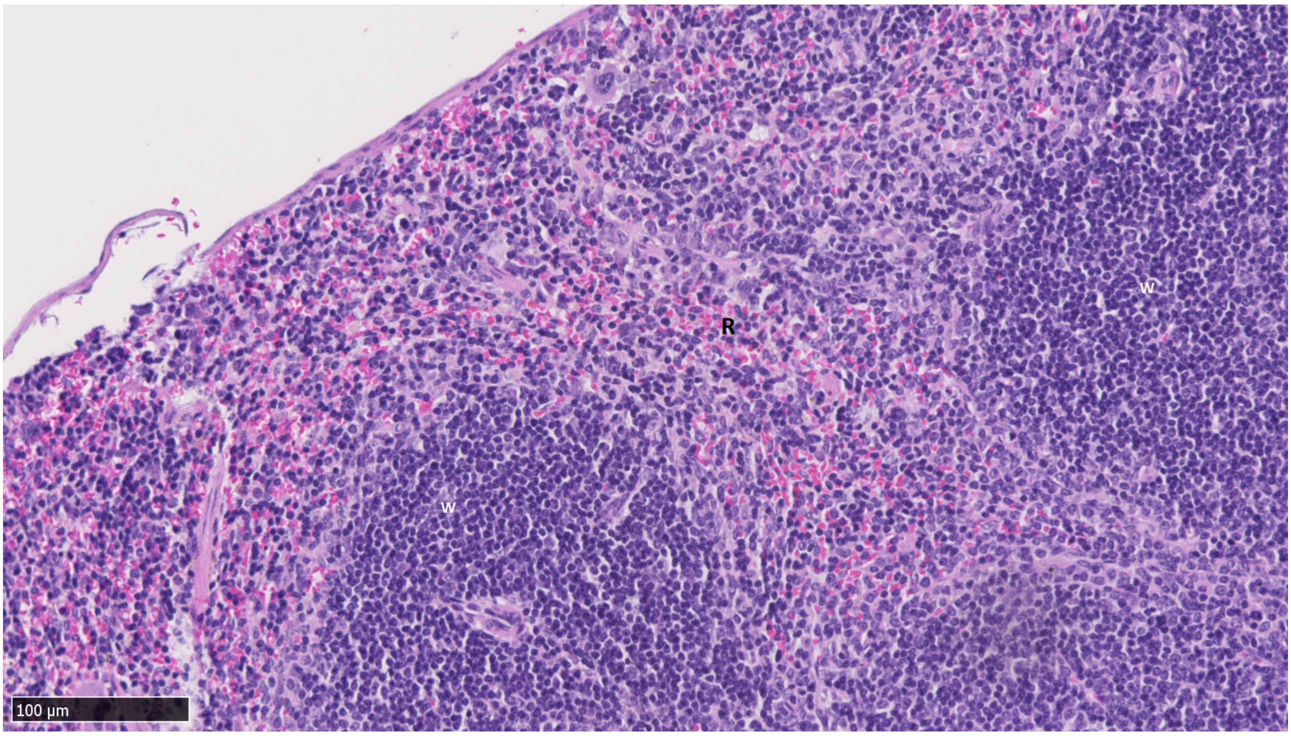

**Supplementary Figure 6:** Spleen close up analysis of mice treated with vehicle. Prominent lymphoid follicles of the white pulp are visible. Compared with treated animals, a mild hematopoiesis in the red pulp is also visible.  W = white pulp, R = red pulp, M = Megakaryocytes.

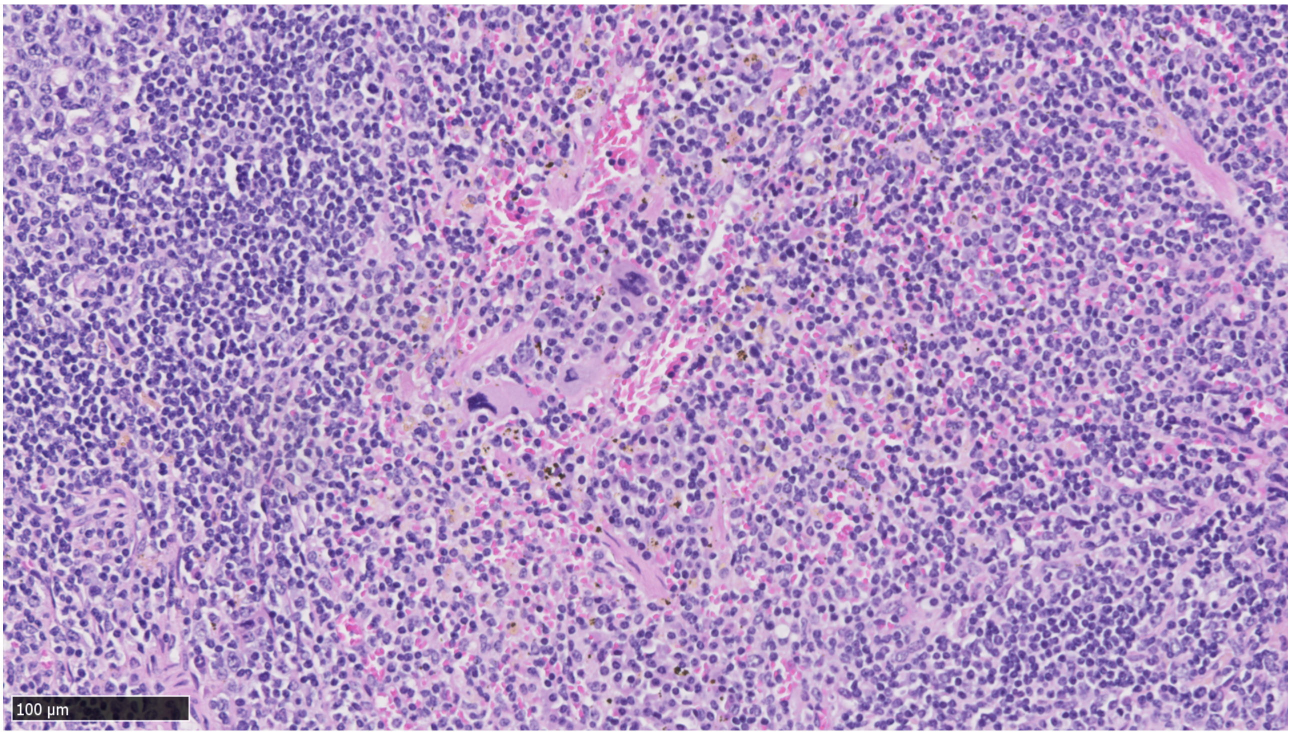

**Supplementary Figure 7:** Spleen close up analysis of mice treated with L19-IL12.

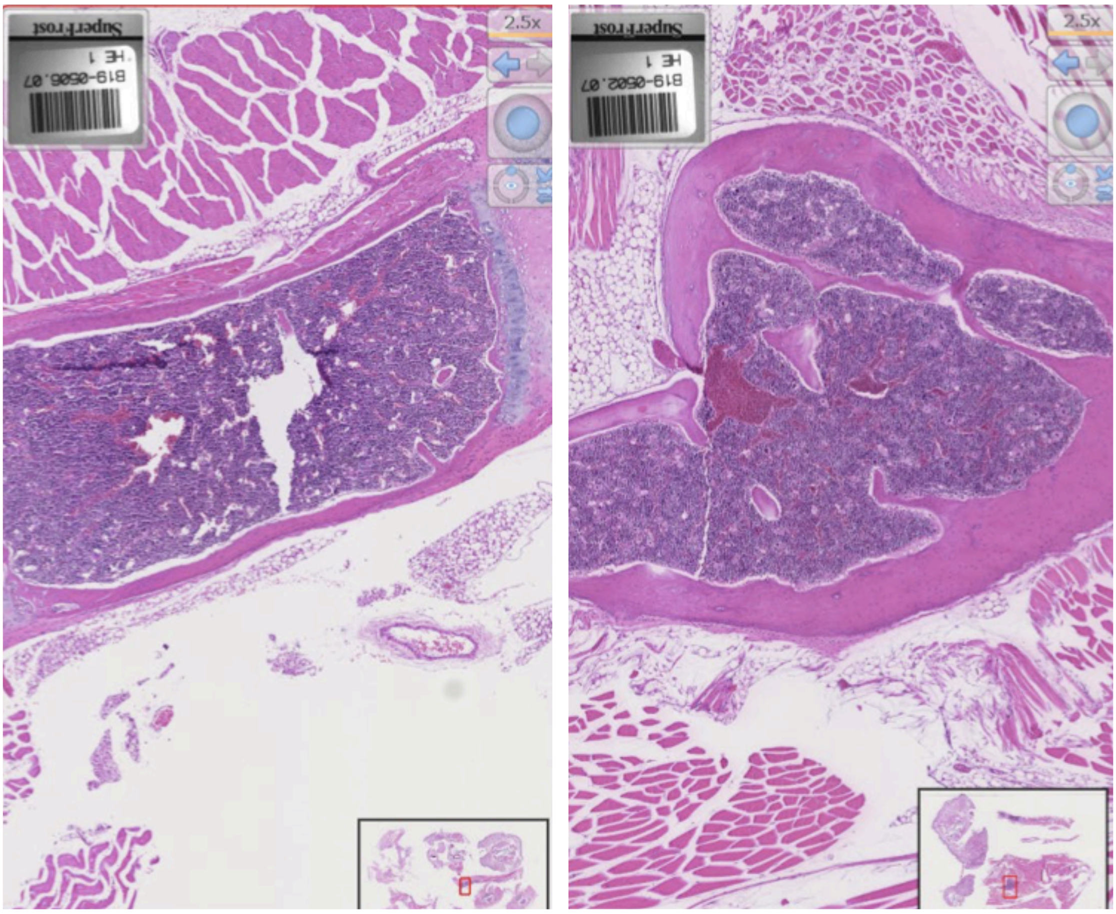

**Supplementary Figure 8:** Bone marrow analysis of mice treated with L19-IL12 (on the left) or with the combination L19-IL12 + AAZ$^{+}$-ValCit-MMAE (on the right)

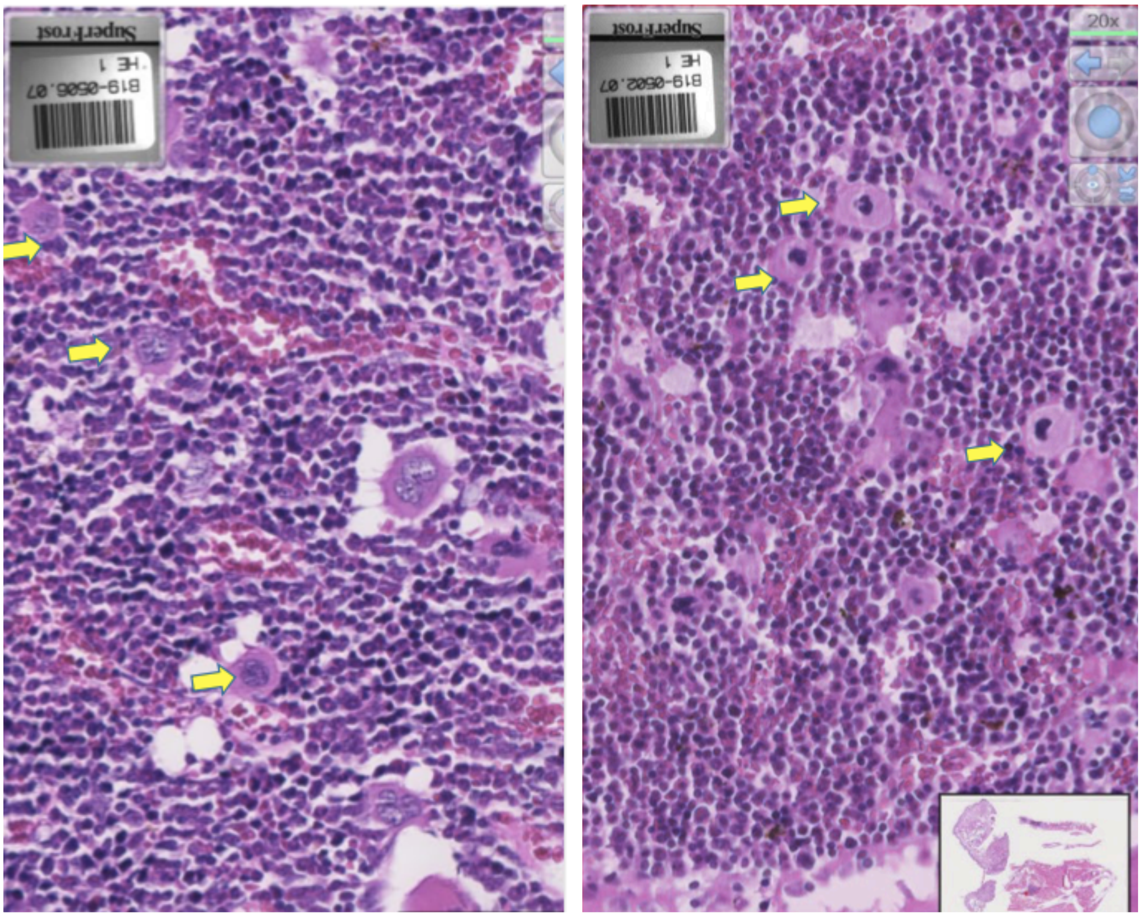

**Supplementary Figure 9:** Bone marrow close-up analysis of mice treated with L19-IL12 (on the left) or with the combination L19-IL12 + AAZ$^{+}$-ValCit-MMAE (on the right). Arrows underline no obvious bone marrow depression.
